# Learning Protein Structure Representations through Joint-Embedding Prediction

**DOI:** 10.64898/2026.11.25.769343

**Authors:** Untaek Hwang, Yoobeen Shin, Jinung Song, Dongwoo Kim, Jaemin Sim, Juyong Lee

**Affiliations:** College of Pharmacy, Seoul National University, 1 Gwanak-ro, Gwanak-gu, Seoul 08826, Republic of Korea; Department of Molecular Medicine and Biopharmaceutical Sciences, Graduate School of Convergence Science and Technology, Seoul National University, Seoul, 08826, Republic of Korea

**Keywords:** protein structure representation learning, self-supervised learning, joint-embedding prediction, backbone reconstruction

## Abstract

**Motivation:** Protein structure representations are often learned by reconstructing coordinates, yet reconstruction quality does not necessarily track performance on biological prediction tasks. We therefore investigate whether joint-embedding prediction can capture information useful for biological prediction while retaining recoverable backbone geometry.

**Results:** We introduce PS-JEPA, a self-supervised model that learns protein structure representations by predicting masked residue embeddings from visible backbone context. The encoder combines geometric graph encoding with bidirectional Mamba and is pretrained without a coordinate-reconstruction objective. On StructTokenBench, frozen PS-JEPA representations achieve a mean remote-homology macro F1 (× 100) of 72.24, compared with 67.12 for GearNet-Edge, the strongest baseline for this category, while remaining competitive in functional and flexibility prediction. A separately trained decoder reconstructs backbones from these frozen representations, achieving a mean C*α* RMSD of 1.61 Å on 309 external test chains filtered to exclude sequence matches to training and reference data. In an exploratory test of unconditional generation, a learned latent prior with the same decoder yields 62 of 384 backbones (16.15%) that meet the computational designability criterion. Together, these results show that joint-embedding prediction can learn representations that support both biological prediction and backbone reconstruction.

**Availability and Implementation:** https://github.com/ahexlwang/PS-JEPA

**Supplementary Information:** Supplementary Methods and Results accompany this manuscript.

## 1. Introduction

Protein structure representation learning aims to encode the biologically relevant information in three-dimensional geometry as features that can be reused across downstream tasks (Zhang et al., 2023; Yuan et al., 2025; Li et al., 2026). Learned representations can support biological tasks such as functional prediction, flexibility prediction, and remote-homology classification, as well as backbone reconstruction and protein generation.

Existing approaches learn structural features through coordinate reconstruction (Li et al., 2026), inverse folding (Dauparas et al., 2022), or contrastive learning (Zhang et al., 2023). However, performance on a pretraining task alone does not establish the biological utility of the learned representations. StructTokenBench (STB) evaluates these three biological tasks, showing that reconstruction quality need not track biological prediction performance (Yuan et al., 2025).

This motivates testing whether prediction in representation space can learn structural features that transfer to biological tasks.

Joint-Embedding Predictive Architecture (JEPA) is a self-supervised framework that predicts target embeddings from observed context (LeCun, 2022). We use this framework to ask whether training without a coordinate-reconstruction loss yields representations that support biological prediction and retain enough geometry for backbone reconstruction.

To this end, we introduce PS-JEPA (Protein Structure Joint-Embedding Predictive Architecture), summarized in Figure 1. The encoder is designed around one property of protein structure: residues that are distant along the chain can be neighbors in three-dimensional space. Starting from node and edge features derived from backbone coordinates (Figure 1(a)), it therefore combines geometric vector perceptrons (GVPs) (Jing et al., 2021b) for local spatial neighborhoods with bidirectional Mamba layers (Gu and Dao, 2024; Liu et al., 2025) for context along the residue chain. During pretraining, we mask contiguous chain segments or spatial neighborhoods and predict embeddings of the masked residues from the visible backbone (Figure 1(b)).

**Figure 1.**
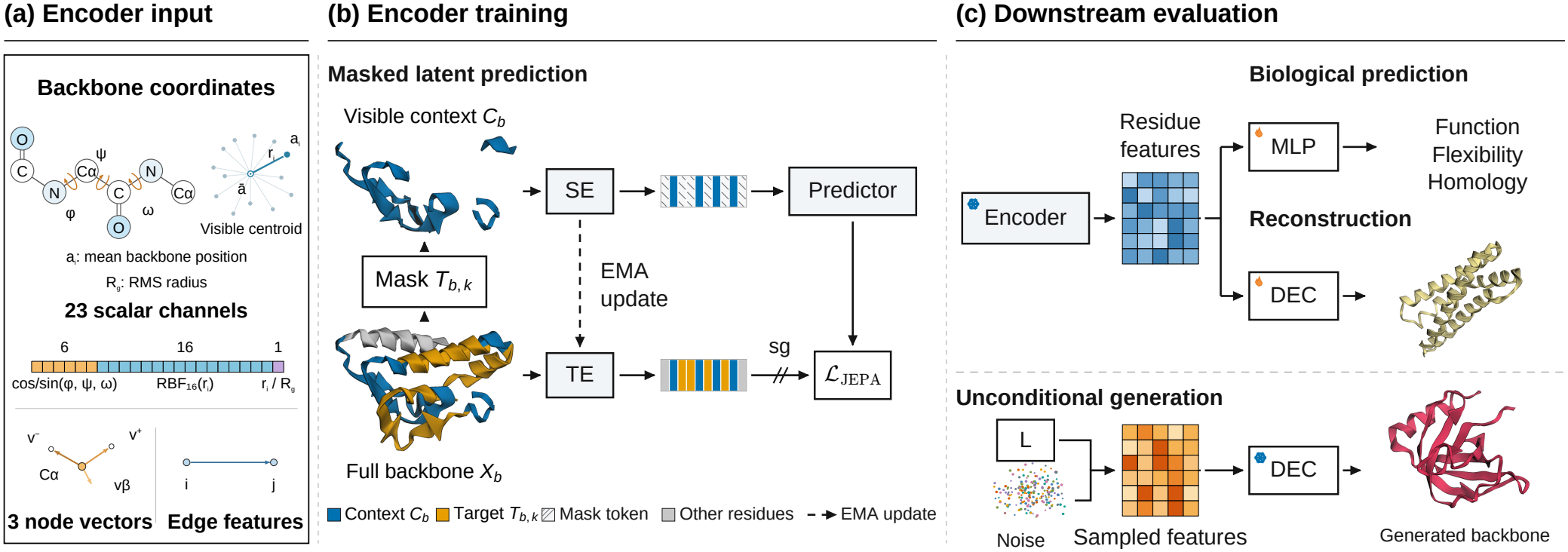
PS-JEPA encoder inputs, pretraining, and downstream evaluation. (a) Backbone coordinates provide 23 scalar and three vector node features, together with edge features. (b) The student encoder (SE) encodes visible context *C*_*b*_ of chain *b*; the predictor estimates teacher embeddings for target block *T*_*b,k*_. The teacher encoder (TE) encodes the full backbone *X*_*b*_ and is updated by an exponential moving average (EMA) of SE weights. The JEPA loss (*L*_JEPA_) uses mean squared error at target positions with stop-gradient (sg) on TE embeddings. Blue and orange mark *C*_*b*_ and *T*_*b,k*_; gray (“Other residues”) marks residues outside *C*_*b*_ ∪ *T*_*b,k*_, and hatching marks mask tokens. (c) Frozen encoder features feed a multilayer perceptron (MLP) for biological prediction or a coordinate decoder (DEC) for backbone reconstruction. For unconditional generation, a learned prior samples residue features from noise conditioned only on chain length *L* (number of residues), and the frozen DEC produces a backbone. Flames denote trainable parameters; snowflakes denote frozen parameters.

We evaluate the biological utility of the learned representations on STB with the encoder frozen (Figure 1(c)). PS-JEPA achieves the highest category-average score among the evaluated models in remote-homology classification, with competitive performance in functional and flexibility prediction. To assess recoverable geometry, we freeze the PS-JEPA encoder and train a separate conditional flow decoder, which reconstructs the backbones of 309 external test chains with a mean C*α* root-mean-square deviation (RMSD) of 1.61 Å. We further explore unconditional backbone generation by learning a prior over residue embeddings and using the same frozen decoder; 62 of 384 generated backbones (16.15%) meet the computational designability criterion after sequence design and refolding.

## 2. Related work

### 2.1. Protein structure representations

Methods for learning protein structure representations differ in their training objectives and in whether they produce discrete codes or continuous features. Foldseek’s 3Di alphabet represents local structure with discrete codes (van Kempen et al., 2024). ESM3’s structure tokenizer (Hayes et al., 2025), ProTokens (Lin et al., 2023a), AminoAseed and its VanillaVQ baseline (Yuan et al., 2025), and Bio2Token (Liu et al., 2025) learn discrete codes through structure reconstruction. MiAE (Chen et al., 2026) and ProteinAE (Li et al., 2026) learn continuous representations through reconstruction.

Inverse folding and contrastive learning are also used to learn structural representations. ProteinMPNN (Dauparas et al., 2022) and MIF (Yang et al., 2023) learn to predict amino-acid sequences conditioned on structure. GearNet introduces a geometric graph encoder together with pretraining methods based on multiview contrastive learning and geometric information prediction (Zhang et al., 2023).

### 2.2. Predictive self-supervised learning

Several self-supervised methods learn representations by predicting or matching learned targets rather than reconstructing inputs. BYOL’s online network takes one augmented view of an image and predicts the embedding that a moving-average target network produces for another view (Grill et al., 2020). DINO learns representations through self-distillation by matching teacher and student output distributions across augmented views (Caron et al., 2021). Given a masked input, data2vec predicts the contextualized representations that its teacher computes from the full input (Baevski et al., 2022).

JEPA frames this idea as predicting target representations from an observed context (LeCun, 2022). I-JEPA applies it to images, using a single context block to predict representations of several target blocks in the same image (Assran et al., 2023). 3D-JEPA and Point-JEPA extend this approach to point clouds by predicting target-region representations from context regions (Hu et al., 2024; Saito et al., 2025).

Recent work has applied joint-embedding prediction to protein sequences and to three-dimensional atomistic systems. ProteinJEPA combines masked amino-acid prediction with latent representation prediction to train protein language models (Ofer et al., 2026). ProtJEPA trains a student encoder that takes sequences as input to predict teacher embeddings integrating multiple biological modalities (Ravideshik et al., 2026). Atom-JEPA performs atom-level and substructure-level latent prediction on three-dimensional molecular and crystalline structures (Petersen et al., 2026). PS-JEPA uses protein backbone coordinates alone to predict an embedding for each masked residue from the visible backbone.

## 3. Methods

PS-JEPA learns protein representations by predicting masked residue embeddings from visible backbone context. We evaluate these representations through three downstream tasks: biological prediction, backbone reconstruction, and unconditional generation (Figure 1(c)). For downstream tasks, we freeze the teacher encoder, whose weights are an exponential moving average (EMA) of the student encoder weights during pretraining. Biological prediction uses task-specific supervised heads, reconstruction uses a coordinate decoder, and unconditional generation combines a learned latent prior with the same frozen decoder. Additional implementation details are provided in Supplementary Sections A.1–A.3.

### 3.1. Datasets

Encoder pretraining and decoder training use the AFDB partitions provided by TEDBench, comprising 742,183 training chains and 7,496 validation chains (Varadi et al., 2022; Chen et al., 2026). The AFDB chains used in this study contain 18–512 residues.

Biological prediction is evaluated on StructTokenBench (STB) using its released task-specific training, validation, and test splits (Yuan et al., 2025).

Reconstruction is evaluated on an external test set of 309 experimental chains from 305 PDB entries (Berman et al., 2000). Candidates are drawn from X-ray structures released between 1 January and 12 November 2026, with chain lengths of 64–512 residues and resolution ≤ 3 Å. We use MMseqs2 (Steinegger and Söding, 2017) to exclude candidates matching AFDB training sequences at ≥ 30% identity and ≥ 80% query coverage.

For unconditional generation, we train the prior on 50,653 filtered AFDB chains of 64–128 residues, with checkpoint selection on 128 fixed validation chains (Supplementary Section D.1). We evaluate 384 generated backbones, 128 each at 80, 100, and 120 residues.

### 3.2. Backbone encoder

The encoder combines local spatial geometry with context along the residue chain. Inputs are N, C*α*, C, and O coordinates (Figure 1(a)). During pretraining, part of the input is masked: residues with withheld backbone coordinates are hidden and the rest are visible (Section 3.3). Each visible residue forms a graph node at the mean position of its backbone atoms and receives messages over directed edges from up to 12 nearest visible neighbors. Three geometric vector perceptron (GVP) blocks (Jing et al., 2021b) encode local geometric features on this directed graph.

Each node carries scalar and vector features. Scalar features are sin and cos of the backbone torsions (*ϕ, ψ, ω*) and encodings of the residue’s distance from the visible-residue centroid. Vector features are unit directions to the next and previous C*α* atoms and a pseudo-C*β* direction derived from the backbone. Edge features encode spatial distance, direction, and signed sequence offset. We use only visible coordinates to build the graph and compute features. We zero torsion channels or C*α* direction vectors that require a hidden atom.

Eight bidirectional Mamba layers (Gu and Dao, 2024) propagate these local geometric features along the residue chain, providing context beyond each spatial neighborhood. Before Mamba processing, hidden residues retain their sequence slots and receive the same learnable mask embedding. Each layer uses shared parameters for its forward and reverse scans and sums their outputs after restoring residue order (Liu et al., 2025). The encoder has approximately 14 million parameters and outputs one 512-dimensional vector per residue. Supplementary Section A.2 details the architecture and features (Figure S1).

### 3.3. Masked latent prediction

For each chain, we choose between masking contiguous chain segments and masking spatial neighborhoods with equal probability, then sample an initial context and four target blocks using the selected mode. All target positions are removed from the initial context, and the four target blocks are predicted separately from the same remaining visible context. Region sizes and sampling procedures are specified in Supplementary Section A.3.

The student encoder *f*_*θ*_ observes the visible context *C*_*b*_ of chain *b*, while the teacher encoder 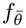 observes the full backbone *X*_*b*_ (Figure 1(b)). The predictor *g*_*ξ*_ consists of two bidirectional Mamba layers and a linear output layer. Each target block is specified by adding a learnable indicator vector to the student encoder output at that block’s positions; predictor parameters are shared across all blocks. For target block *T*_*b,k*_, the prediction at residue *i* is 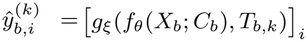. Targets are 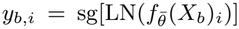, where non-affine LayerNorm standardizes channels within each residue and sg stops gradients. Let *T*_*k*_ contain the pairs (*b, i*) in the microbatch for which *i* is a non-padding position in *T*_*b,k*_. We minimize

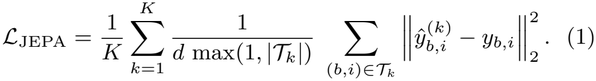

The loss averages over *d* = 512 channels and valid target positions within each block, then weights the *K* = 4 blocks equally; empty blocks contribute zero. The student encoder and predictor are trained with this loss, while the teacher encoder is updated by an EMA of the student encoder’s parameters.

#### Training settings

Optimization uses AdamW, with learning-rate reductions and stopping based on validation JEPA loss. For the main STB comparison, we use the teacher encoder from each of three independent runs (seeds 42–44) at its validation-based stopping point. For reconstruction and generation, we continue the seed-42 run to 35,000 updates and freeze the resulting teacher encoder. Full training settings and the update at which each run stopped are given in Supplementary Section A.3.

### 3.4. Downstream evaluation

#### 3.4.1. Biological prediction

STB (Yuan et al., 2025) evaluates frozen structural representations through task-specific supervised heads (Figure 1(c)).

##### Baseline methods

Figure 2 shows 14 representations from 13 methods. PS-JEPA provides continuous features and their post-hoc vector quantization with 512 codes. The 12 external baselines are Foldseek 3Di, ProTokens, the ESM3 structure tokenizer, VanillaVQ, AminoAseed, Bio2Token, ProteinMPNN, MIF, MiAE-S, MiAE-B, GearNet-Edge, and ProteinAE (Section 2). We use released weights or tokenizers and train only the supervised heads. Supplementary Table S7 shows published STB scores in parentheses. The compared methods differ in representation format (discrete or continuous), input modality, training data, and objectives. MIF and GearNet-Edge additionally use amino-acid identities, whereas PS-JEPA uses only backbone coordinates.

**Figure 2.**
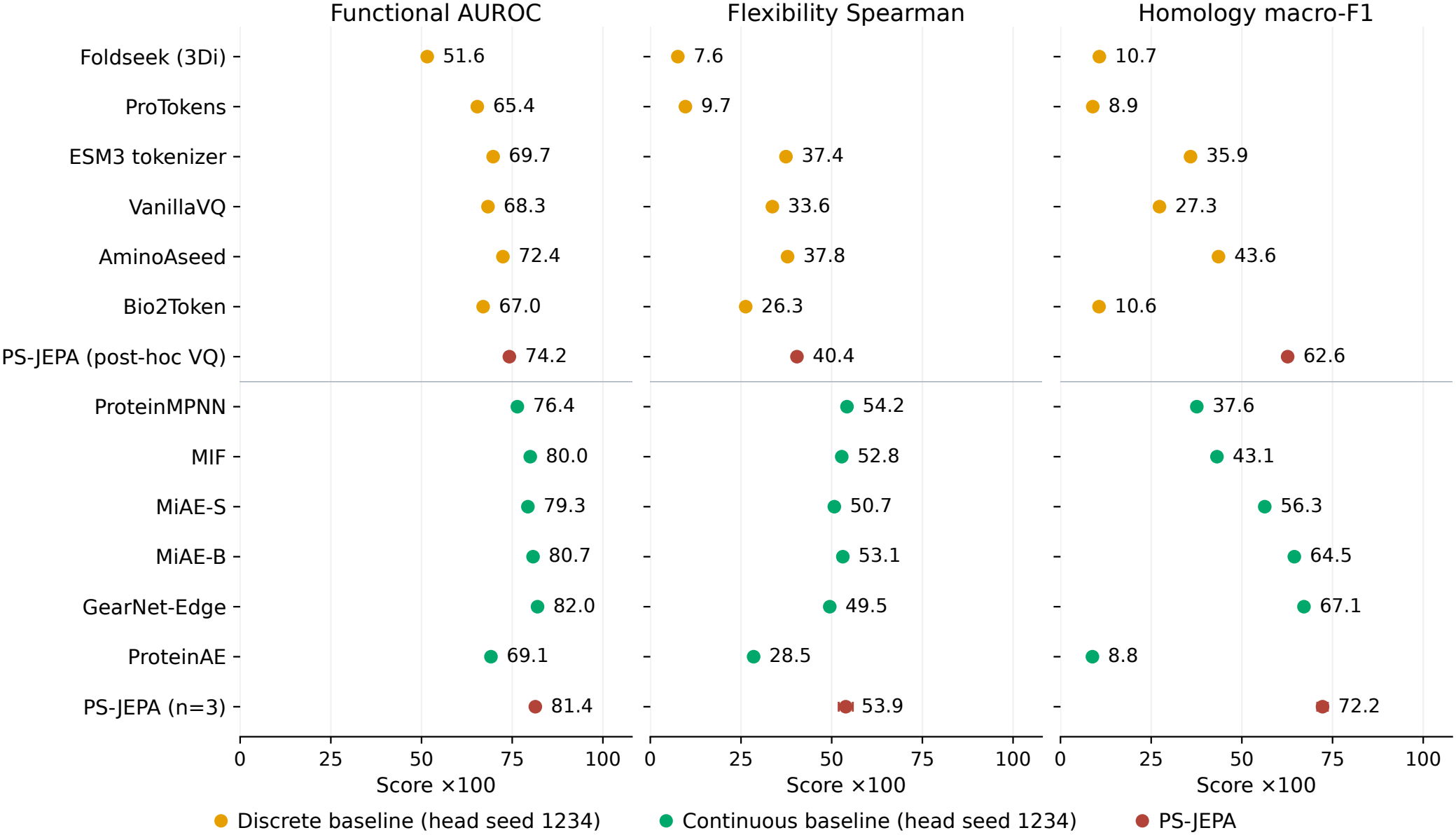
Biological prediction from 14 frozen representations on StructTokenBench (STB). Panels average functional AUROC (15 conditions), flexibility Spearman correlation (six), and homology macro F1 (three) on original STB labels (*×*100, higher is better). Discrete (orange) and continuous (green) baselines use head seed 1234. Both PS-JEPA rows (red) average head seeds 1234, 2345, and 3456 per encoder. The discrete PS-JEPA row uses Post-hoc VQ (512 codes) with one 35,000-update encoder (seed 42; Table S6(a)). Continuous PS-JEPA points and bars show the mean *±*1 sample SD across encoder seeds 42–44. Table S7 details the other 13 representations.

#### Pretraining and initialization controls

We compare PS-JEPA with two controls. Coordinate-reconstruction encoders assess the choice of pretraining protocol, and randomly initialized encoders assess the contribution of pretraining. We train reconstruction encoders jointly with attention decoders for 35,000 updates (independent seeds 42–44), then freeze the encoders for STB evaluation. The random encoders use the PS-JEPA inputs and architecture without pretraining and remain frozen (seeds 42–44). For both controls we average three heads per encoder and report the mean and sample SD across encoders. The reconstruction loss and architecture are detailed in Supplementary Section A.4.4.

##### Heads and metrics

With each encoder frozen, we train task-specific heads on the training splits, select learning rates and checkpoints on the validation splits, and report final scores on the test splits. Functional-site and flexibility heads predict residue-level labels. Homology classification uses mean pooling over valid residue features to obtain a chain representation. The main comparison uses the original released STB labels. The masking-policy (Figure S2D), architecture (Table S5), and chain-mixing (Table S6(b)) controls use labels corrected for mapping and target-precision errors (corrected-v1, Supplementary Section B.1). Each control uses one label version. PS-JEPA averages head seeds 1234, 2345, and 3456 per encoder. Continuous PS-JEPA reports mean ± sample SD across three encoders, while post-hoc VQ uses encoder seed 42. In the main comparison, each external baseline uses head seed 1234.

Pairing each of the 12 STB tasks with its test splits gives 24 evaluation conditions. Each category score is the mean over the conditions in that category: 15 functional AUROCs, six flexibility Spearman correlations, and three homology macro F1 scores. Following the released STB implementation, functional AUROC and homology macro F1 are computed per batch and averaged without weighting. Flexibility Spearman correlation is computed from valid predictions and targets concatenated across the evaluation set. Head architectures, checkpoint selection, and metric aggregation are detailed in Supplementary Section B.1.

#### 3.4.2. Backbone reconstruction

We freeze the 35,000-update seed-42 PS-JEPA encoder (Section 3.3) and train a separate conditional flow decoder (Lipman et al., 2023), as illustrated in Figure 1(c). The decoder takes noisy backbone coordinates, the flow time, and the 512-dimensional residue embeddings of the full input backbone, and predicts clean backbone coordinates. Its output head predicts a rigid frame and a carbonyl rotation for each residue, from which the N, C*α*, C, and O coordinates are constructed.

Decoder training proceeds from initial coordinate fitting to global-shape refinement and then fine-tuning on intermediate states generated by the decoder itself. Reconstruction starts from independent Gaussian noise and uses 100 Euler steps.

The external test set, encoder and decoder checkpoints, and inference settings are fixed before evaluation. We report one reconstruction for every test chain. Aligned C*α* RMSD measures coordinate recovery; reference-normalized TM-score (Zhang and Skolnick, 2004) and the C*α* local distance difference test (lDDT) (Mariani et al., 2013) measure overall fold and local distance agreement, respectively. The decoder architecture, training losses and schedules, and evaluation details are given in Supplementary Section C.1.

#### 3.4.3. Unconditional generation

We train a flow-based latent prior over residue embeddings extracted by the frozen teacher encoder, *Z* = *E*(*X*) ∈ ℝ^*L×*512^, where 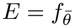. The embeddings are standardized channelwise, *Y* = (*Z* − *µ*)*/σ*, where *µ* and *σ* are the channel means and standard deviations computed from encoder embeddings of the prior training chains. The prior takes noisy features, the chain length, and the flow time, and predicts the clean features *Y*. It is trained with a time-weighted mean squared error between the predicted and clean features and an auxiliary geometry loss, which compares backbones decoded from its predictions with the training backbones; the gradients of this loss pass through the frozen decoder and update only the prior.

Generation is conditioned only on chain length *L*, without a specified amino-acid sequence or structural motif. The prior transforms *L* Gaussian noise vectors into features, which we pass to the frozen reconstruction decoder after reversing the feature standardization (Figure 1(c)).

We generate backbones with four sampling seeds, 32 per length and seed. For each backbone, ProteinMPNN designs eight sequences, which are refolded with ESMFold (Dauparas et al., 2022; Lin et al., 2023b). A backbone meets the computational designability criterion if at least one refold has full-chain C*α* self-consistency RMSD (scRMSD) below 2 Å (Geffner et al., 2025). All 384 backbones are included in the denominator, without ranking or filtering; a backbone is counted as not designable if none of its eight sequence-design and refolding attempts yields a structure satisfying this criterion. Further settings are given in Supplementary Section D.

## 4. Results

### 4.1. Biological prediction

#### 4.1.1. StructTokenBench comparison

PS-JEPA achieves the highest homology category score among the 14 evaluated representations, with competitive performance in functional and flexibility prediction (Figure 2). Its homology category score is 72.24, exceeding the highest-scoring baseline, GearNet-Edge (67.12), by 5.12 points.

The advantage is particularly pronounced on the Fold split, which tests fold recognition on proteins from superfamilies absent from the downstream training set (Yuan et al., 2025). On this split, PS-JEPA scores 55.61, exceeding the best baseline score of 46.62 from MiAE-B by 8.99 points (Supplementary Table S7). PS-JEPA also scores higher than GearNet-Edge, MiAE-B, and MiAE-S in the homology category in sensitivity analyses using alternative F1 aggregation rules and additional head seeds (Supplementary Table S8).

For functional and flexibility prediction, PS-JEPA scores 81.38 and 53.88, respectively, slightly below the best baseline scores of 81.98 (GearNet-Edge) and 54.22 (ProteinMPNN). The discrete PS-JEPA representation (post-hoc VQ with 512 codes, one 35,000-update encoder) scores higher than all six discrete baselines in each category, with the largest margin in homology (62.62 versus 43.56 for the best discrete baseline, AminoAseed; Figure 2 and Supplementary Table S6(a)).

#### 4.1.2. Encoder controls

##### Pretraining and initialization

PS-JEPA and coordinate-reconstruction encoders both outperform randomly initialized encoders across all three STB categories (Table 1). The largest difference between PS-JEPA and coordinate reconstruction is in homology, where PS-JEPA scores 20.33 points higher; the corresponding differences in functional and flexibility prediction are 1.68 and 0.44 points.

**Table 1.** Pretraining and initialization on original STB labels.

| Encoder | Functional | Flexibility | Homology |
|---|---|---|---|
| PS-JEPA | 81.38 ± 0.14 | 53.88 ± 1.94 | 72.24 ± 1.45 |
| Coordinate reconstruction | 79.70 ± 0.10 | 53.44 ± 0.52 | 51.91 ± 1.56 |
| Random | 74.85 ± 0.13 | 44.27 ± 0.12 | 27.14 ± 0.63 |

Category means: AUROC over 15 functional conditions, Spearman correlation over six flexibility conditions, and macro F1 over three homology conditions. PS-JEPA uses EMA teacher encoders; reconstruction uses non-EMA encoders trained for 35,000 updates. Each row averages head seeds 1234, 2345 and 3456 within each encoder, then reports the mean ± sample SD across independent encoder seeds 42–44 (*n* = 3). Protocol details are given in Supplementary Section A.4.4.

##### Chain-context processing

When the encoder’s bidirectional Mamba layers are removed and the GVP blocks are expanded to match the parameter count, homology F1 falls from 72.65 to 62.20, while functional and flexibility scores exceed the reference (82.97 versus 81.80 and 56.51 versus 54.27; Supplementary Table S6(b)).

Without this expansion, homology F1 falls further to 52.04 and functional prediction drops below the reference (79.89). This pattern supports a contribution of chain-context processing to homology prediction beyond total parameter count; it does not indicate an advantage across all tasks. These controls use 35,000-update encoders and corrected-v1 labels, averaging three heads per encoder. The reference is the mean of three encoders (seeds 42–44), whereas each ablation is a single encoder (seed 42).

### 4.2. Backbone reconstruction

A decoder conditioned on frozen PS-JEPA representations reconstructs the 309 external test chains with a mean C*α* RMSD of 1.61 Å (Table 2; distributions in Figure 3). Mean TM-score and C*α*-lDDT are 0.938 and 0.877, respectively.

**Table 2.**
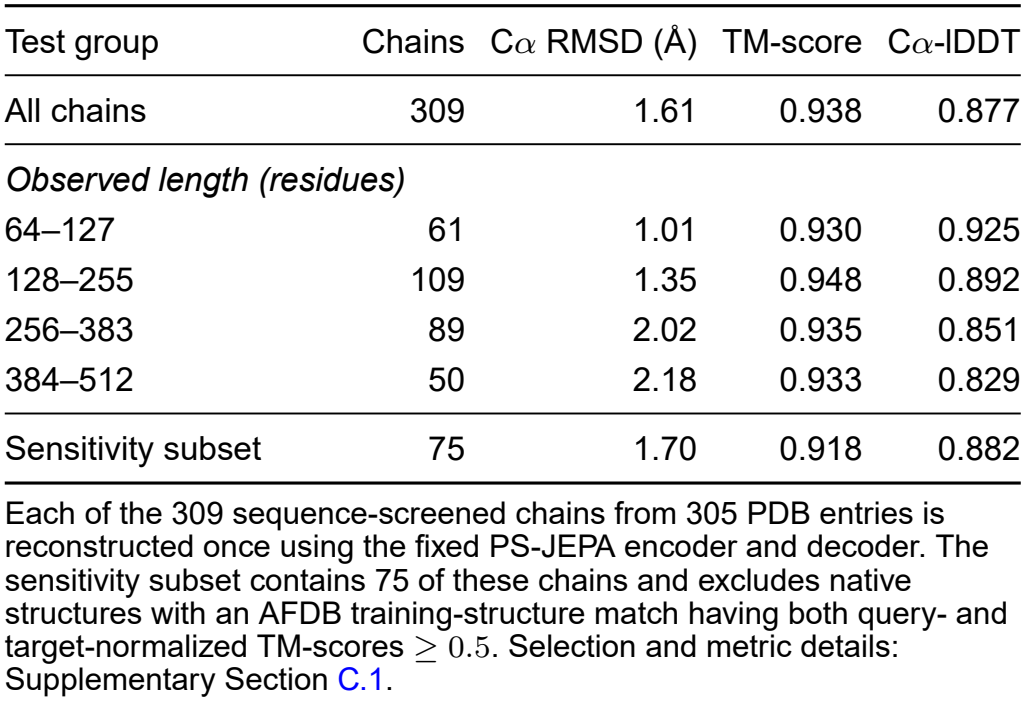
Backbone reconstruction on the external test set.

**Figure 3.**
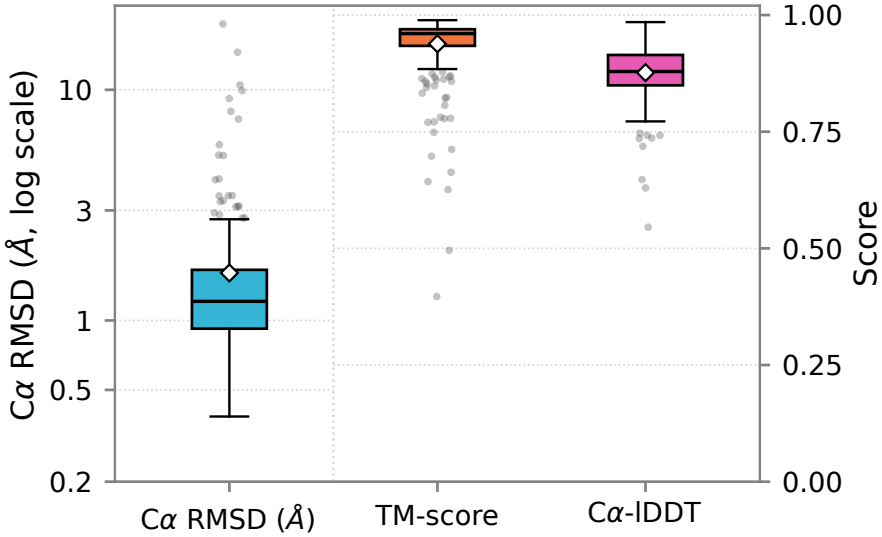
Reconstruction distributions on the external test. Results include all 309 experimental chains of 64–512 observed residues. Boxes show interquartile ranges (IQRs), lines indicate medians, and diamonds indicate means. Whiskers extend to the most extreme values within 1.5 IQR of the box; all outliers are shown. Aligned C*α* RMSD uses the left logarithmic axis; TM-score and C*α*-lDDT use the right linear axis.

Mean C*α* RMSD increases with chain length (Table 2). Figure 4 shows reconstructions near the median C*α* RMSD for three length groups. For the sensitivity analysis, we exclude chains with a match to an AFDB training structure for which both the query- and target-normalized TM-scores are at least For the remaining 75 chains, mean C*α* RMSD is 1.70 Å, mean TM-score is 0.918, and mean C*α*-lDDT is 0.882, close to the corresponding means for the full test set.

**Figure 4.**
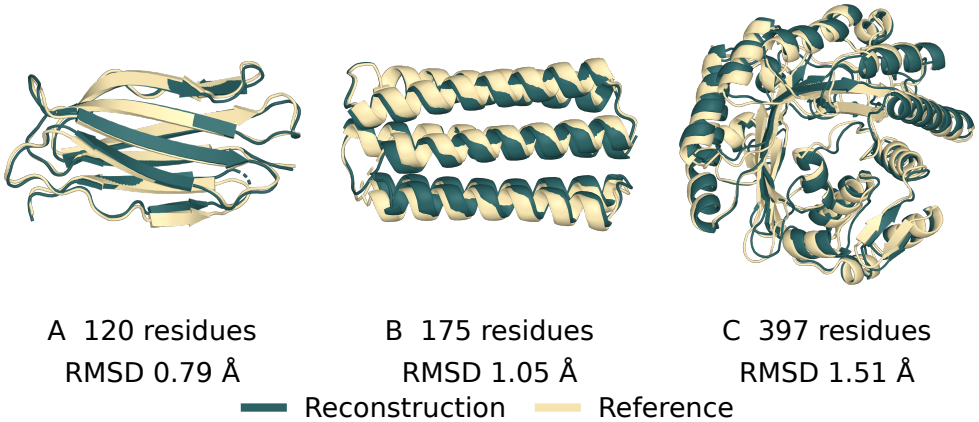
(A) 11OS chain A, (B) 9R2B chain B, and (C) 9P3C chain A. Cases have C*α* RMSDs near the medians of the 64–127, 128–255, and 256–512 residue groups, respectively.

### 4.3. Unconditional generation

With the learned latent prior and the frozen reconstruction decoder, 62 of 384 generated backbones pass the computational designability criterion (16.15%; Table 3). Designability is highest at 80 residues (25.78%) and lower at 100 and 120 residues (10.94% and 11.72%, respectively). Figure 5 shows designable examples. Generation and evaluation protocols are described in Supplementary Section D.

**Table 3.** Designability rates of unconditionally generated backbones.

| Length | Generated | Designable | Designability (%) |
|---|---|---|---|
| 80 | 128 | 33 | 25.78 |
| 100 | 128 | 14 | 10.94 |
| 120 | 128 | 15 | 11.72 |
| All | 384 | 62 | 16.15 |

For each backbone, eight sequences are designed with ProteinMPNN and refolded with ESMFold. A backbone is classified as designable if at least one refold has full-chain Cα self-consistency RMSD (scRMSD) below 2 Å relative to the generated backbone.

**Figure 5.**
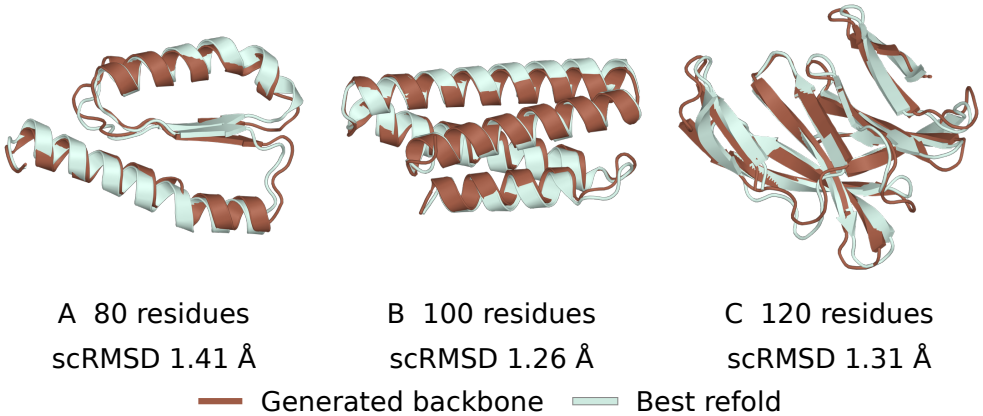
Selected designable backbones from unconditional backbone generation. Generated backbones are aligned to their best ESMFold refolds among eight ProteinMPNN-designed sequences, based on full-chain C*α* scRMSD. One selected designable backbone is shown at each length.

## 5. Discussion and conclusion

We explore how to adapt JEPA to protein backbones by modeling both the order of residues along the chain and their spatial relationships in the folded structure. JEPA models for point clouds operate on raw coordinates (Hu et al., 2024; Saito et al., 2025), but a protein has no canonical orientation, and its features should not change when the structure is rotated. We therefore encode each residue’s spatial neighborhood with GVP blocks (Jing et al., 2021b), whose scalar outputs are unchanged by rotation. These blocks capture local geometry, but a fold is a property of the whole chain, so we add bidirectional Mamba layers (Gu and Dao, 2024) to integrate the local features along the residue chain. Both GVP encoders without Mamba score below the reference encoder on homology, including the parameter-matched encoder (Supplementary Table S6(b)), which suggests that this chain-level integration aids remote-homology classification.

The ablations affect the three task categories differently. Longer training and greater Mamba depth raise homology scores without corresponding gains in flexibility (Supplementary Tables S4 and S5). When Mamba is removed and the GVP blocks are expanded to match the parameter count, the encoder scores above the reference encoder in both functional and flexibility prediction (Supplementary Table S6(b)). Functional and flexibility prediction may therefore depend more on local geometric features than on their integration along the residue chain.

PS-JEPA’s homology performance suggests that representations reflecting global structural relationships in proteins can be learned without training to reconstruct coordinates (Figure 2; Table 1). Predicting full-structure features from partial observations may encourage the student encoder to learn relationships between visible and masked regions (Assran et al., 2023).

The representations also retain backbone geometry. Although the encoder is never trained to reconstruct coordinates, a separately trained decoder recovers the backbones of 309 external test chains from frozen PS-JEPA representations with a mean C*α* RMSD of 1.61 Å (Table 2). Features learned by joint-embedding prediction therefore support biological prediction while retaining enough geometry for backbone reconstruction.

This study has several limitations. The AFDB pretraining set was not filtered against the STB evaluation proteins, so proteins identical or homologous to them may have been seen during pretraining. Pretraining uses backbone coordinates only and never accesses STB labels, so this exposure cannot leak labels. It could still favor homology classification if folds seen during pretraining are easier to separate. An evaluation split that removes proteins similar to the pretraining set would show how much of the homology performance depends on such exposure.

Generation remains limited. With the same decoder, only 62 of 384 backbones generated from sampled features meet the computational designability criterion (Table 3). Because the encoder learns masked feature prediction and the prior is fitted separately on frozen representations, it remains unclear which stage limits designability. Structural novelty of the generated backbones relative to the training set of the prior was not assessed.

PS-JEPA takes only backbone coordinates as input and could be extended by incorporating complementary information such as the amino-acid sequence. The representation also remains informative after discretization: when quantized post hoc to 512 codes, it scores above all six discrete baselines in every STB category (Figure 2; Supplementary Table S6(a)). Such discrete structure tokens could be paired with protein language models.

In conclusion, PS-JEPA shows that joint-embedding prediction, with no coordinate-reconstruction objective, can learn protein structure representations that support both biological prediction and backbone reconstruction. Future work could improve generation quality and pursue the extensions outlined above.

## Author Contributions

Untaek Hwang designed the study, developed the method, implemented the optimization framework, performed the experiments, and analyzed the results.

Yoobeen Shin, Jinung Song, Dongwoo Kim, and Jaemin Sim contributed to conceptualization, manuscript writing, and analysis.

Juyong Lee supervised the project and contributed to study design, conceptualization, and manuscript revision.

All authors reviewed and approved the final manuscript.

## Supplementary Material

Supplementary Methods and Results are included in this manuscript.

## Data and Code Availability

The datasets used in this study are available from AFDB/TEDBench, StructTokenBench, and the Protein Data Bank (Varadi et al., 2022; Chen et al., 2026; Yuan et al., 2025; Berman et al., 2000). The PS-JEPA code and pretrained checkpoints are available at https://github.com/ahexlwang/PS-JEPA.

## Supplementary Methods and Results

### A. Encoder details

This appendix describes datasets, the encoder, and PS-JEPA pretraining (Supplementary Sections A.1–A.3). Ablations and design controls are grouped in Supplementary Section A.4.

#### A.1. Data and optimization

We use the AFDB/TEDBench training partition for encoder pretraining and decoder fitting. Biological prediction uses the released task-specific STB splits. Table S1 summarizes the data partitions and optimization settings.

**Table S1.**
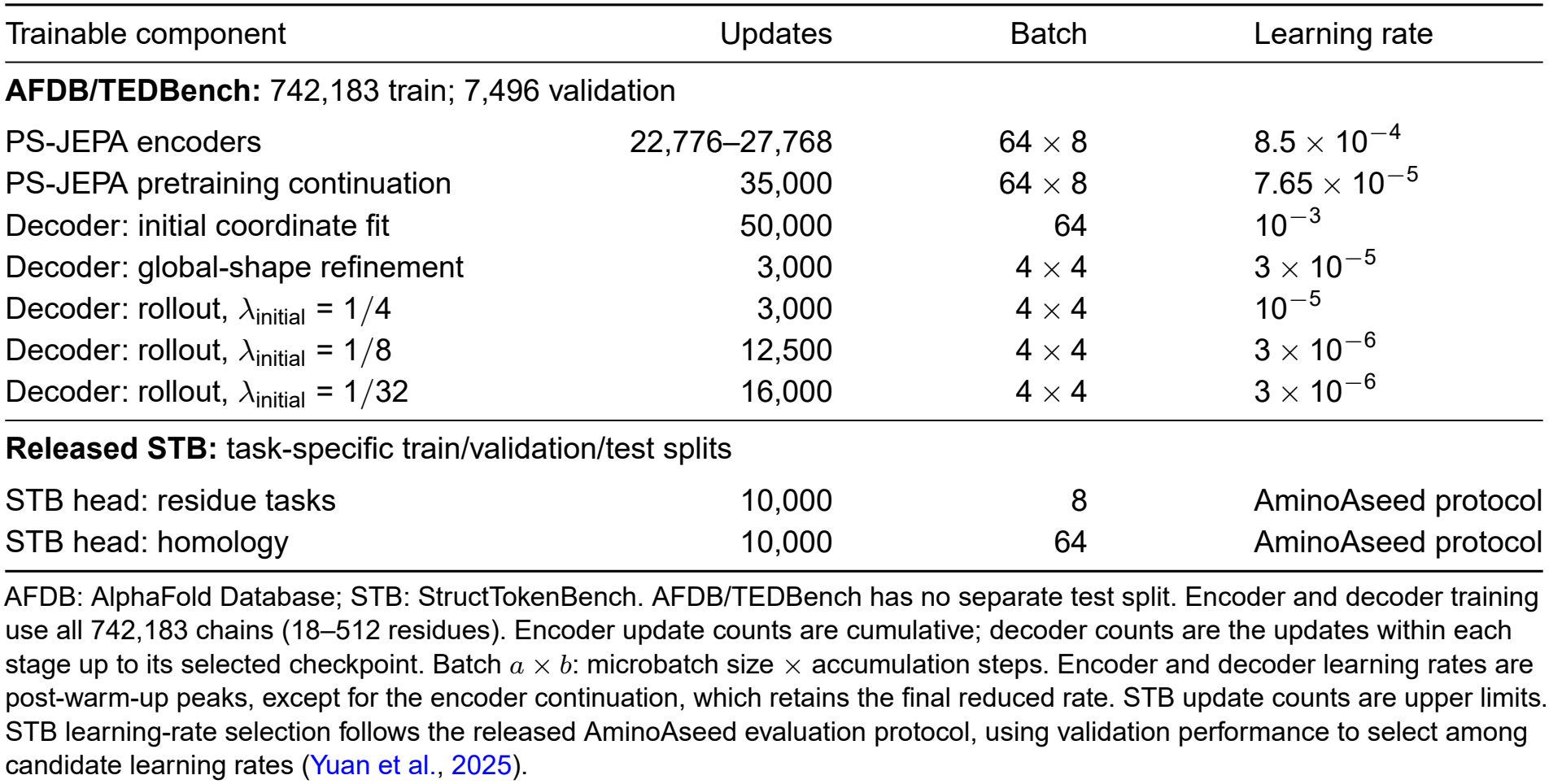
Data partitions and training settings.

| Trainable component | Updates | Batch | Learning rate |
|---|---|---|---|
| **AFDB/TEDBench:** 742,183 train; 7,496 validation |  |  |  |
| PS-JEPA encoders | 22,776–27,768 | 64 × 8 | 8.5 × 10<sup>−4</sup> |
| PS-JEPA pretraining continuation | 35,000 | 64 × 8 | 7.65 × 10<sup>−5</sup> |
| Decoder: initial coordinate fit | 50,000 | 64 | 10<sup>−3</sup> |
| Decoder: global-shape refinement | 3,000 | 4 × 4 | 3 × 10<sup>−5</sup> |
| Decoder: rollout, *λ*<sub>initial</sub> = 1/4 | 3,000 | 4 × 4 | 10<sup>−5</sup> |
| Decoder: rollout, *λ*<sub>initial</sub> = 1/8 | 12,500 | 4 × 4 | 3 × 10<sup>−6</sup> |
| Decoder: rollout, *λ*<sub>initial</sub> = 1/32 | 16,000 | 4 × 4 | 3 × 10<sup>−6</sup> |
| **Released STB:** task-specific train/validation/test splits |  |  |  |
| STB head: residue tasks | 10,000 | 8 | AminoAseed protocol |
| STB head: homology | 10,000 | 64 | AminoAseed protocol |

AFDB: AlphaFold Database; STB: StructTokenBench. AFDB/TEDBench has no separate test split. Encoder and decoder training use all 742,183 chains (18–512 residues). Encoder update counts are cumulative; decoder counts are the updates within each stage up to its selected checkpoint. Batch *a* × *b*: microbatch size × accumulation steps. Encoder and decoder learning rates are post-warm-up peaks, except for the encoder continuation, which retains the final reduced rate. STB update counts are upper limits. STB learning-rate selection follows the released AminoAseed evaluation protocol, using validation performance to select among candidate learning rates (Yuan et al., 2025).

#### A.2. Geometric encoder

The encoder combines geometric message passing over visible residues with bidirectional processing along the residue chain (Figure S1). Three GVP blocks encode spatial neighborhoods, and eight bidirectional Mamba layers integrate the resulting features in residue order. Hidden positions retain their sequence slots and receive the same learnable mask embedding before chain processing.

##### Inputs and graph

Coordinates *X* ∈ ℝ^*L×*4*×*3^ contain N, C*α*, C, and O. The atom mask *A*_*ia*_ marks available atoms; *R* denotes valid (non-padding) residues and *C* ⊆ *R* the visible context. The teacher encoder uses the full view, *C* = *R*; the student encoder uses the context view. The TEDBench (Chen et al., 2026) AFDB source provides N/C*α*/C; O is reconstructed from local peptide geometry using the next residue’s N when available, with a local-frame fallback at chain ends.

We subtract the mean available C*α* position and apply one random rigid rotation per chain during training. Validation uses the same centering without rotation. Each chain is zero-padded at the end to the longest chain length in its batch; atom and residue masks mark padding as invalid.

The graph nodes *V* are visible residues with at least one available atom. We represent residue *i* by the mean position *a*_*i*_ of its available backbone atoms. Each node receives messages from up to the 12 nearest other nodes, using distances between these mean positions, without adding reverse edges. Edges and their features are computed once per view and reused across the three GVP blocks.

**Figure S1.**
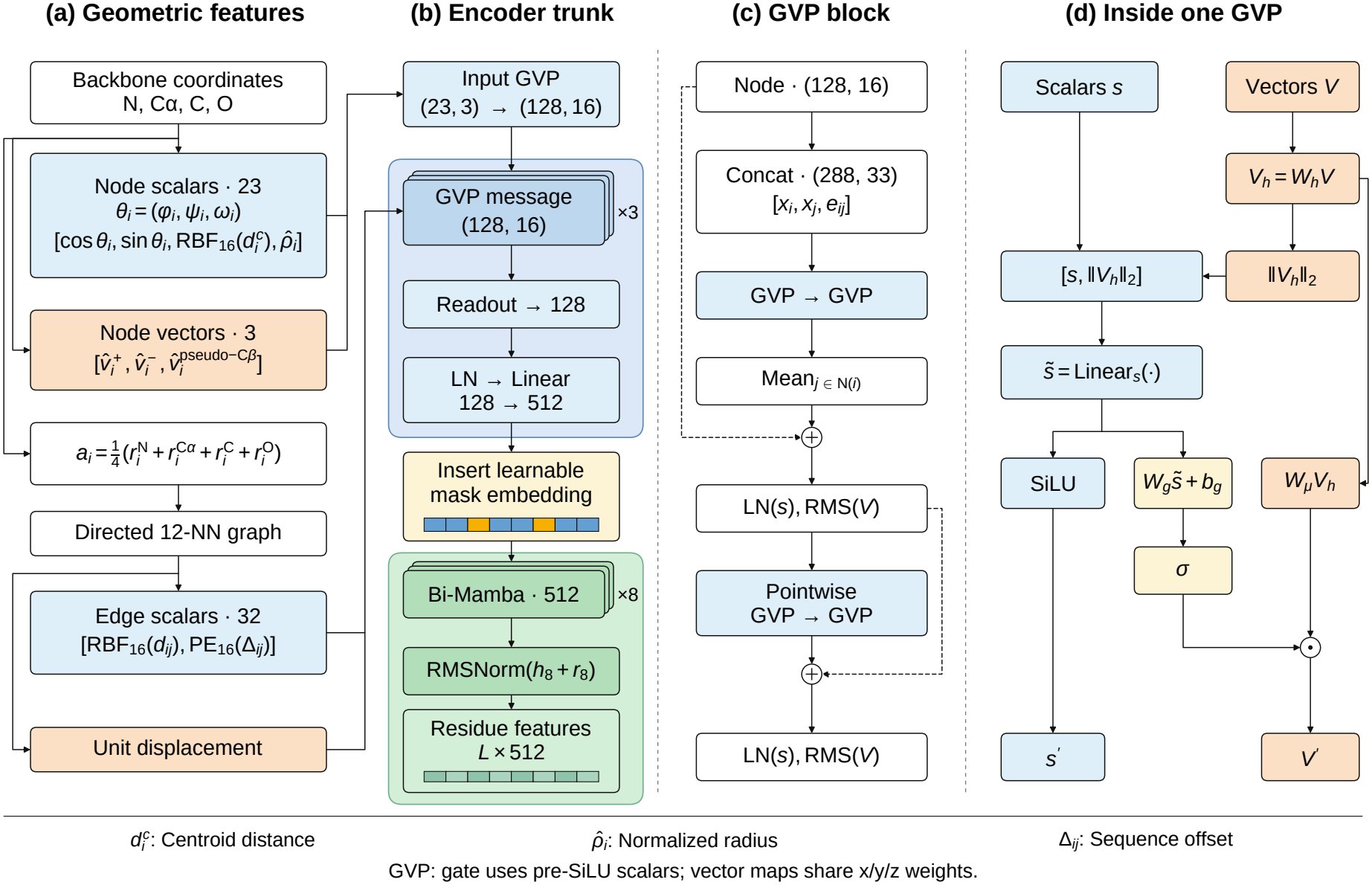
PS-JEPA encoder architecture. (a) Backbone-derived node and edge features; the mean backbone position formula assumes all four atoms are present. In (a), 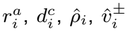 and Δ_*ij*_ correspond to *x*_*i,a*_, *r*_*i*_, *r*_*i*_*/R*_*g*_, the forward and backward C*α* directions, and the sequence offset *j − i* in Table S2. The pseudo-C*β* direction 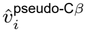 corresponds to 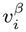, and the edge distances satisfy *d*_*ij*_ = *d*_*ji*_. (b) Three GVP blocks encode local geometry, and eight bidirectional Mamba layers integrate context along the residue chain. Hidden positions receive the same learnable embedding. Blue/yellow slots are visible/hidden residues; green cells depict residue features. (c) GVP message and pointwise updates; (*s, v*) counts scalar and three-component vector channels. (d) Internal scalar and vector transformations within a gated GVP. RBF: radial basis functions; PE: positional encoding; NN: nearest neighbors; LN: layer normalization; RMS: root-mean-square normalization; Bi-Mamba: bidirectional Mamba. Cell diagrams are schematic.

##### Geometric features

Node features describe local backbone geometry and distance from the visible-residue centroid. Edge features describe spatial separation, direction, and sequence offsets between neighboring residues. All distances are in Å. Table S2 defines the channels using backbone torsions (*ϕ, ψ, ω*) and normalized directions *u*(*v*) = *v/* max(∥*v*∥_2_, 10^*−*8^). The pseudo-C*β* direction is 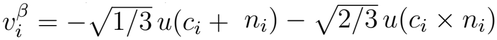, where *n*_*i*_ = *u*(*x*_*i,N*_ *− x*_*i,Cα*_) and *c*_*i*_ = *u*(*x*_*i,C*_ *− x*_*i,Cα*_). Radial channels use *r*_*i*_ = ∥*a*_*i*_ − *ā*∥_2_, where *ā* averages the mean backbone positions *a*_*i*_ over visible residues and *R*_*g*_ is their root-mean-square distance to *ā*. Context and teacher views compute these statistics separately.

**Table S2.**
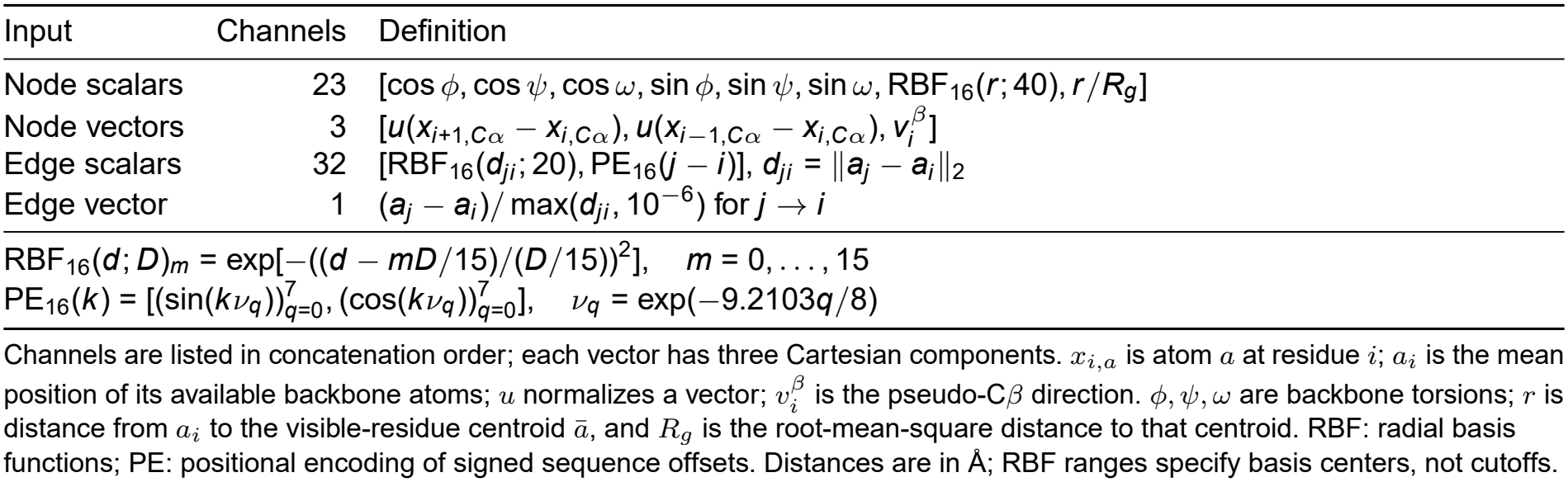
Geometric input channels of the backbone encoder.

| Input | Channels | Definition |
|---|---|---|
| Node scalars | 23 | [cos ϕ, cos ψ, cos ω, sin ϕ, sin ψ, sin ω, RBF<sub>16</sub>(*r*; 40), *r*/*R<sub>g</sub>*] |
| Node vectors | 3 | [*u*(*x*<sub>*i*+1,*C*α</sub> − *x*<sub>*i*,*C*α</sub>), *u*(*x*<sub>*i*−1,*C*α</sub> − *x*<sub>*i*,*C*α</sub>), *v*<sub>*i*</sub><sup>β</sup>] |
| Edge scalars | 32 | [RBF<sub>16</sub>(*d<sub>ji</sub>*; 20), PE<sub>16</sub>(*j* − *i*)], *d<sub>ji</sub>* = ‖***a***<sub>*j*</sub> − ***a***<sub>*i*</sub>‖<sub>2</sub> |
| Edge vector | 1 | (***a***<sub>*j*</sub> − ***a***<sub>*i*</sub>)/ max(*d<sub>ji</sub>*, 10<sup>−6</sup>) for *j* → *i* |
| RBF<sub>16</sub>(*d*; *D*)<sub>*m*</sub> = exp[−((*d* − *mD*/15)/(*D*/15))<sup>2</sup>], *m* = 0, . . . , 15 |  |  |
| PE<sub>16</sub>(*k*) = [(sin(*kν<sub>q</sub>*))<sup>7</sup><sub>*q*=0</sub>, (cos(*kν<sub>q</sub>*))<sup>7</sup><sub>*q*=0</sub>], ν<sub>*q*</sub> = exp(−9.2103*q*/8) |  |  |

Channels are listed in concatenation order; each vector has three Cartesian components. *x*<sub>*i*,*a*</sub> is atom *a* at residue *i*; *a*<sub>*i*</sub> is the mean position of its available backbone atoms; *u* normalizes a vector; *v*<sub>*i*</sub><sup>β</sup> is the pseudo-Cβ direction. ϕ, ψ, ω are backbone torsions; *r* is distance from *a*<sub>*i*</sub> to the visible-residue centroid ā, and *R*<sub>*g*</sub> is the root-mean-square distance to that centroid. RBF: radial basis functions; PE: positional encoding of signed sequence offsets. Distances are in Å; RBF ranges specify basis centers, not cutoffs.

##### Missing coordinates, masking, and sequence boundaries

For torsions formed within the input array, both sine and cosine channels are set to zero if any of the four defining atoms is missing or hidden. Forward and backward C*α* directions require visible C*α* atoms at both residues; the pseudo-C*β* direction requires visible N, C*α*, and C atoms at the same residue. Otherwise, the corresponding direction vector is set to 0. Undefined angles at the input-array ends are instead padded with the angle-0 encoding, (cos, sin) = (1, 0).

Sequence neighbors are adjacent entries in the input array. Hidden residues retain their slots; sequence neighbors are not reassigned to skip hidden positions. Entries that are consecutive in the array remain sequence neighbors even when their C–N distance is large.

##### GVP and chain updates

We use gated GVPs (Jing et al., 2021a,b) with (128, 16) scalar–vector hidden channels, an input mapping from (23, 3), and three residual blocks. Each block contains a two-GVP message network with mean aggregation over valid neighbors, followed by a two-GVP pointwise network. Messages concatenate receiver, sender, and edge features; empty neighborhoods contribute zero. Both updates use SiLU scalar activations, sigmoid vector gates, and scalar–vector normalization (LayerNorm (Ba et al., 2016) *ϵ* = 10^*−*5^; vector mean-square floor 10^*−*8^). A scalar-only GVP readout, LayerNorm, and a 128 → 512 projection supply the chain features.

The same learnable 512-dimensional mask vector replaces projected features at hidden residue slots; the full-view teacher and downstream encoding use projected features at every valid slot. The chain model has eight width-512 bidirectional Mamba layers (Gu and Dao, 2024), with state size 16, convolution width 4, and expansion 2. Each layer shares forward and reverse parameters and, after restoring residue order, sums the directional outputs separately for the layer-output (*h*) and residual (*r*) streams.

Scans include hidden slots; trailing padding stays in place and is zeroed before each layer and at the output. We use RMSNorm with *ϵ* = 10^*−*5^ (Zhang and Sennrich, 2019) and FP32 residual accumulation, normalize the final sum of the two streams, and omit absolute position embeddings.

The resulting encoder output contains one 512-dimensional representation per valid residue, with padded positions set to zero.

#### A.3. Pretraining

Pretraining learns residue representations by predicting full-view teacher embeddings at target positions from the visible context.

##### Context and target construction

Sequence-contiguous masking selects a contiguous interval in the ordered valid-residue list; spatial masking selects the nearest residues around a C*α* seed. Each chain chooses either mode with probability 0.5 and uses it for the initial context and four independently sampled target blocks. For *n* valid residues, we set the sizes of the initial context and each target block to max(1, round(*cn*)) and max(1, round(*τ n*)), respectively, with *c* = 0.85 and *τ* = 0.125. Chains with fewer than four valid residues receive no context or targets. Spatial seeds and neighboring residues are selected from valid residues with an available C*α* atom. Let *C*_0_ denote the initial context and *T*_*k*_ the *k*th target block. The visible context is

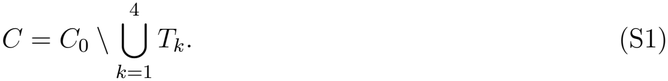

The 85% context fraction is applied before target removal, so it is not the final visible fraction. All four target blocks are predicted from the same remaining context. Target blocks may overlap. Residues outside the initial context that belong to no target block are hidden from the student encoder and receive no direct prediction loss.

##### Block-conditioned prediction

Each target block is specified by adding a learnable 512-dimensional indicator vector to the student encoder embeddings at that block’s residue positions. The predictor uses two width-512 bidirectional Mamba layers and a 512 × 512 linear projection with bias. It processes the full valid sequence, but the loss uses only the target positions. All four predictions share predictor parameters and the same student encoder output.

##### Teacher targets and loss reduction

Each teacher residue embedding *u* ∈ ℝ^512^ is standardized across channels using non-affine LayerNorm:

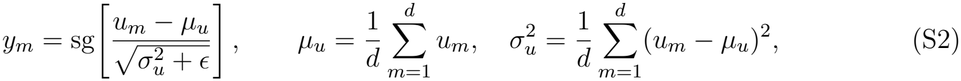

where *m* indexes channels, *d* = 512, *ϵ* = 10^*−*5^, and sg blocks gradient propagation. Normalization acts within each residue, without learned scale or offset; predictor outputs are not normalized.

The sole objective, Equation (1), averages squared errors over channels and valid target residues across each microbatch, then equally across the four blocks. Residues shared by multiple target blocks contribute to the loss for each corresponding block; empty blocks contribute zero. Each optimizer update accumulates eight microbatches of 64 chains:

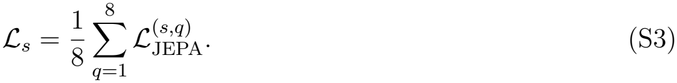

##### Teacher updates

The teacher encoder is initialized from the student encoder and updated once after each optimizer step, using the newly updated student encoder parameters:

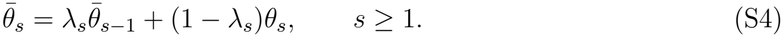

Here *θ*_*s*_ denotes the student encoder parameters after update *s*. The EMA momentum after *s* optimizer updates is

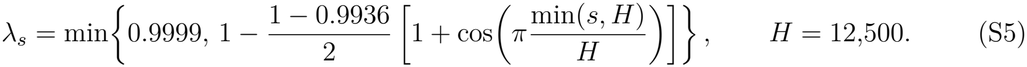

The EMA momentum increases from 0.9936 and is capped at 0.9999.

##### Optimizer and learning-rate schedule

AdamW (Loshchilov and Hutter, 2019) jointly updates the student encoder and predictor, using (*β*_1_, *β*_2_) = (0.9, 0.95), *ϵ*_AdamW_ = 10^*−*8^, and a weight decay of 0.05. The accumulated gradient norm is clipped to 1 before each update. Training uses BF16 mixed precision, with linear warm-up followed by the validation-triggered plateau schedule in Table S3. Encoder training uses seeds 42–44.

##### Validation and plateau criterion

Validation runs every 312 optimizer updates on 50 fixed batches of 64 chains. The plateau criterion is met when the mean JEPA loss over the three most recent evaluations fails to improve on the preceding best value by more than 0.5% for six consecutive checks. Each learning-rate reduction resets the averaging window, best value, and patience counter. At most two reductions are applied before stopping, and reductions or stopping are permitted from update 12,500. An evaluation is flagged for representation collapse if the effective rank of the teacher targets is below 8 or their mean channel standard deviation is below 0.05, with both statistics averaged over validation batches. Flagged evaluations are excluded from the plateau calculation.

**Table S3.** Principal-encoder learning-rate schedule.

| Phase | Condition | Learning rate |
|---|---|---|
| Linear warm-up | First 625 updates | (8.5 × 10<sup>−4</sup>) *s*/625 |
| Peak learning rate | After warm-up, before the first reduction | 8.5 × 10<sup>−4</sup> |
| First reduction | Plateau criterion met | 2.55 × 10<sup>−4</sup> |
| Second reduction | Plateau criterion met again | 7.65 × 10<sup>−5</sup> |
| Stopping | Plateau criterion met after two reductions | — |

*s* is the optimizer-update number, starting at one. Each reduction multiplies the learning rate by 0.3 and applies from the next update.

##### Principal checkpoints and continuation

The principal STB evaluations use the final checkpoints from the validation-based stopping rule: updates 27,768, 22,776, and 23,400 for seeds 42, 43, and 44, respectively. We continued the seed-42 run from 27,768 to 35,000 updates at its final learning rate (7.65 × 10^*−*5^), retaining the EMA schedule and disabling plateau-based stopping. Reconstruction, generation, and post-hoc quantization use the teacher encoder at 35,000 updates.

##### Ablation schedules

Matched ablations trained from scratch use the separate cosine schedule in Supplementary Section A.4.3. For the graph-density comparison in Supplementary Section A.4.4, both runs are evaluated at 42,000 updates, shortened from the planned 60,000 endpoint before STB evaluation without changing the learning-rate schedule.

### A.4. Ablations and controls

#### A.4.1. Masking

We vary the initial context fraction *c* and per-block target fraction *τ* under mixed masking (Figure S2A–C). These sweeps use width-128 encoders with four Mamba layers, width-128 heads, an effective pretraining batch size of 512, a peak learning rate of 1.2 × 10^*−*3^, the plateau criterion described in Supplementary Section A.3, and the original-label validation protocol.

The principal encoder uses *c* = 0.85 and *τ* = 0.125, which balance functional and FlexRMSF scores against homology (Figure S2A–C). At this setting with *K* = 4, an audit of 32 validation chains with five draws per policy finds no visible–target overlap, and all three policies compared in Figure S2D retain approximately half of the residues as visible context (47.22% for sequence-only, 49.44% for spatial-only, and 48.39% for mixed masking).

**Figure S2.**
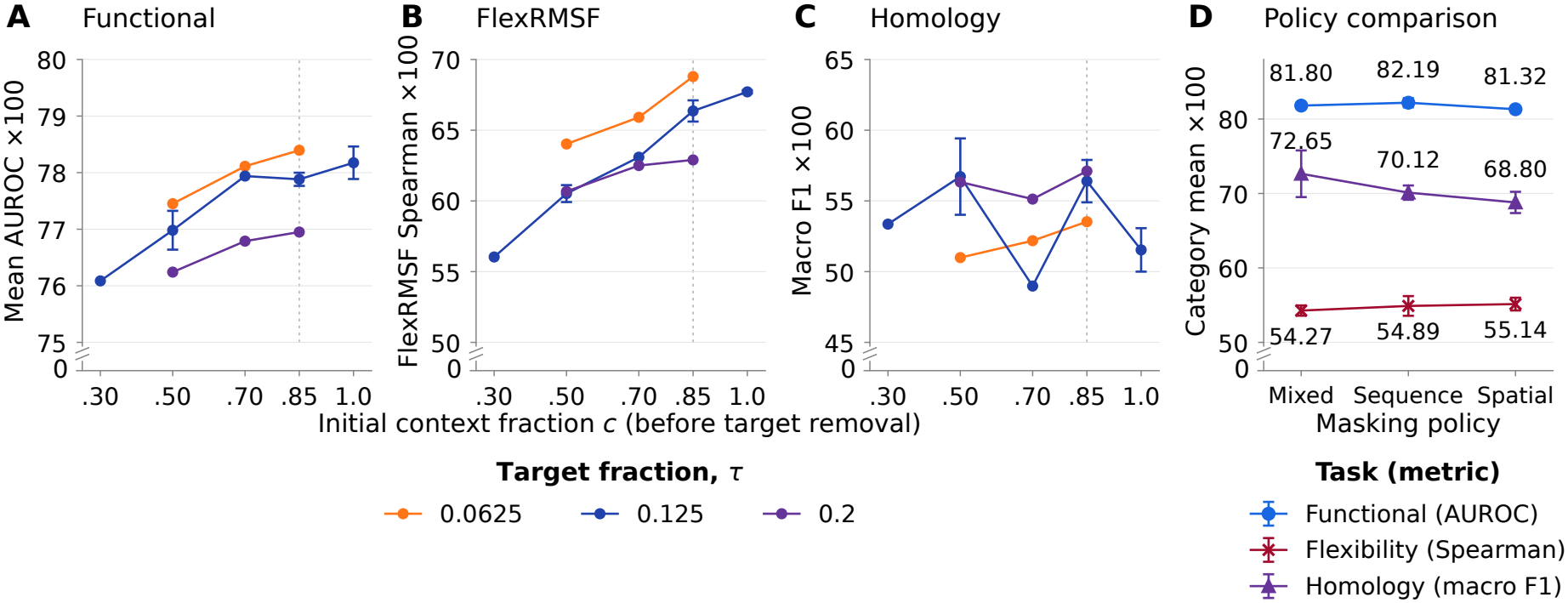
Masking-ratio and policy ablations. (A–C) Width-128 masking sweeps on original-label validation data: (A) mean AUROC over eight functional tasks, (B) FlexRMSF Spearman correlation, and (C) homology macro F1. Points show means and bars show min–max ranges over encoder seeds 42 and 43 where both are available; otherwise, only seed 42 is shown. Vertical dashed lines in A–C mark *c* = 0.85, the initial context fraction used by the principal encoder. (D) Width-512 masking policies at 35,000 updates on corrected-label test data: mean *±* sample SD over encoder seeds 42–44 (*n* = 3), after averaging head seeds 1234, 2345, and 3456 within each encoder. All scores are multiplied by 100; broken axes use panel-specific linear scales.

#### A.4.2. Training duration

We retrospectively evaluate frozen EMA checkpoints to examine how pretraining duration affects downstream utility. A separate seed-42 run replays the recorded principal-encoder learning-rate schedule to a fixed 35,000-update endpoint. It differs from the plateau-continuation encoder used in Table S6(a). We evaluate checkpoints every 1,000 updates from 5,000 to 35,000 on original-label StructTokenBench, independently fitting heads with seeds 1234, 2345, and 3456 at each checkpoint. Table S4 reports selected checkpoints at 5,000-update intervals. Here, ± denotes the sample SD across the three head seeds for one fixed encoder checkpoint, rather than across independently pretrained encoders.

The effect of longer pretraining differs across tasks (Table S4; Figure S3). Homology macro F1 (multiplied by 100) increases from 66.17 ± 0.69 at 5,000 updates to 76.13 ± 1.14 at 35,000, with smaller net gains after approximately 15,000 updates and fluctuations between later checkpoints. Functional scores remain near 81–82, whereas flexibility decreases from 58.15±0.25 to 55.35±0.42. Additional pretraining therefore favors homology in this run without uniformly improving all downstream categories.

**Table S4.** StructTokenBench performance across pretraining updates.

| Updates | Func. | Flex. | Homo. |
|---|---|---|---|
| 5k | 81.34 ± 0.37 | 58.15 ± 0.25 | 66.17 ± 0.69 |
| 10k | 81.96 ± 0.08 | 56.13 ± 0.41 | 71.23 ± 2.58 |
| 15k | 81.71 ± 0.32 | 55.98 ± 0.63 | 75.17 ± 3.46 |
| 20k | 81.72 ± 0.14 | 55.77 ± 0.55 | 75.40 ± 1.23 |
| 25k | 81.55 ± 0.16 | 55.63 ± 0.46 | 76.59 ± 1.55 |
| 30k | 81.43 ± 0.14 | 55.56 ± 0.28 | 74.49 ± 1.62 |
| 35k | 81.24 ± 0.18 | 55.35 ± 0.42 | 76.13 ± 1.14 |

Mean ± sample SD over head seeds (*n* = 3); encoder fixed. Func.: functional AUROC; Flex.: flexibility Spearman correlation; Homo.: homology macro F1. All scores are multiplied by 100; k = 1,000 updates.

**Figure S3.**
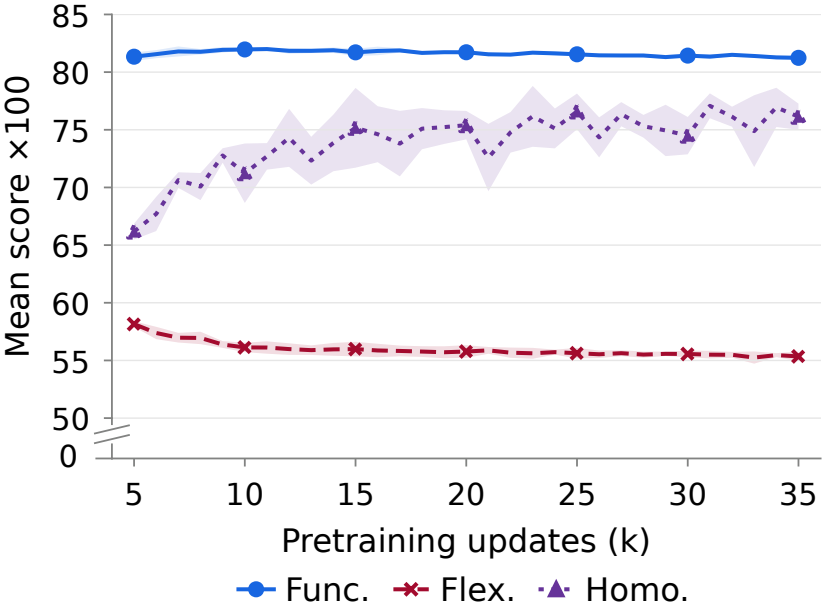
Effect of pretraining duration. Longer pretraining improves homology prediction, with most gains occurring within the first 15,000 updates. Functional performance remains stable, while flexibility prediction declines. Curves show means and shaded bands indicate *±*1 SD across three head seeds at each checkpoint of a single encoder training run.

#### A.4.3. Encoder architectures

To assess how architectural choices affect STB performance, we vary the depths of the GVP, Mamba encoder, and predictor and compare eight-layer Mamba and Transformer chain mixers (Table S5). We pretrain these variants separately from the principal encoder, using a model width of 512 and a cosine learning-rate schedule for 35,000 updates. One- and five-layer GVP controls fix the Mamba encoder at eight layers and the predictor at two.

##### LeJEPA-style control

The LeJEPA-style control (Balestriero and LeCun, 2025) uses the same GVP–Mamba architecture with an eight-layer Mamba encoder. It combines multi-view invariance with sketched isotropic Gaussian regularization (SIGReg) at both protein and residue levels, regularizing projected embeddings toward an isotropic Gaussian distribution. It uses no predictor or EMA teacher; projectors are discarded for downstream evaluation.

##### Architecture effects

With three GVP layers and a two-layer predictor, homology F1 rises with Mamba depth, from 70.86 with four layers to 75.59 with eight and 76.31 with twelve, while encoder parameters increase from 7.20M to 20.76M. With one and five GVP layers (encoder seed 42), homology F1 is 70.74 and 74.84, respectively: 4.85 and 0.75 below the three-layer reference. Their functional/flexibility scores are 81.59/54.91 and 81.78/54.20. Increasing predictor depth does not uniformly improve performance: four layers improve functional and flexibility scores but reduce homology F1 relative to two layers. The eight-layer Transformer gives much lower homology F1 than the eight-layer Mamba reference (41.45 versus 75.59) despite having more parameters (25.90M versus 13.98M). These comparisons use equal update counts rather than equal computational cost, as model sizes differ.

**Table S5.** StructTokenBench architecture ablations.

| Model | GVP | Chain mixer | Pred. | Enc. (M) | Func. | Flex. | Homo. |
|---|---|---|---|---|---|---|---|
| PS-JEPA | 3 | Mamba 4 | 2 | 7.20 | 81.61 | 57.71 | 70.86 |
| PS-JEPA | 3 | Mamba 8 | 1 | 13.98 | 81.73 | 52.12 | 71.30 |
| PS-JEPA | 3 | Mamba 8 | 2 | 13.98 | 81.53 | 54.74 | 75.59 |
| PS-JEPA | 3 | Mamba 8 | 4 | 13.98 | 82.78 | 57.63 | 71.47 |
| PS-JEPA | 3 | Mamba 12 | 2 | 20.76 | 81.86 | 54.57 | 76.31 |
| PS-JEPA | 3 | Transformer 8 | 2 | 25.90 | 79.40 | 51.46 | 41.45 |
| LeJEPA-style | 3 | Mamba 8 | — | 13.98 | 72.95 | 37.98 | 55.53 |
| PS-JEPA | 1 | Mamba 8 | 2 | 13.76 | 81.59 | 54.91 | 70.74 |
| PS-JEPA | 5 | Mamba 8 | 2 | 14.20 | 81.78 | 54.20 | 74.84 |

Corrected-v1; width 512; 35,000 updates; encoder seed 42 and head seed 1234 for all rows (*n* = 1). GVP, mixer, Pred.: layer counts; Enc. (M): encoder parameters in millions, excluding predictor. Func./Flex./Homo.: mean functional AUROC/flexibility Spearman/homology macro F1 (×100). Transformer: eight heads, feed-forward expansion 4, no dropout, learned positions.

##### Training-recipe comparison

The LeJEPA-style control scores below the matched PS-JEPA reference in all three categories, including 55.53 versus 75.59 in homology (Table S5). The comparison concerns the full training recipes: view construction, predictor use, and EMA targets differ alongside the regularization objective.

#### A.4.4. Encoder controls

Table S6 examines representation format, encoder chain mixing, and graph construction. Table 1 in the main text compares pretraining methods and random initialization. Encoder weights are frozen during head training, and all evaluations cover the same 24 STB conditions. Training budgets, label versions, and input geometry are specified for each control. Tables 1 and S6 average the three head seeds within each encoder, with the independent encoder count specified for each comparison.

**Table S6.** Encoder controls.

| Configuration | *n* | Functional | Flexibility | Homology |
|---|---|---|---|---|
| *(a) PS-JEPA encoder: 35k updates; original labels* |  |  |  |  |
| Continuous | 1 | 81.32 | 56.03 | 72.65 |
| Post-hoc VQ (512 codes) | 1 | 74.22 | 40.42 | 62.62 |
| FSQ-4096 | 1 | 74.15 | 43.76 | 28.40 |
| *(b) Chain mixing: 35k updates; corrected-v1 labels* |  |  |  |  |
| GVP–Bi-Mamba reference | 3 | 81.80 ± 0.22 | 54.27 ± 0.69 | 72.65 ± 3.13 |
| Without encoder Bi-Mamba layers | 1 | 79.89 | 55.09 | 52.04 |
| Without Bi-Mamba, parameter matched | 1 | 82.97 | 56.51 | 62.20 |
| *(c) Graph construction: original labels* |  |  |  |  |
| *k* = 12 (42k updates) | 1 | 81.10 | 53.63 | 72.76 |
| *k* = 30 (42k updates) | 1 | 81.95 | 55.90 | 72.83 |
| Cα-position graph (35k) | 3 | 81.35 ± 0.29 | 53.46 ± 0.96 | 72.93 ± 1.61 |

Category means (×100) cover 15 functional AUROCs, six flexibility Spearman correlations and three homology macro F1s. Head seeds 1234, 2345 and 3456 are averaged within each encoder. Figure 2 uses the same three-head mean for the Post-hoc VQ (512 codes) row. *n* counts independent encoders: seed 42 for *n* = 1, seeds 42–44 for *n* = 3 (mean ± sample SD). The mean-position reference in (a) uses plateau continuation; Table S4 uses a separate learning-rate replay. VQ/FSQ numbers give codebook sizes; *k* gives the number of graph neighbors.

##### Post-hoc quantization

We freeze the PS-JEPA encoder at 35,000 updates and quantize its representations post hoc. VQ assigns each residue embedding, standardized with statistics from AFDB training chains, to the nearest of 512 *k*-means centers. FSQ-4096 (Mentzer et al., 2024) projects to six scalar channels, quantizes each to four levels, and projects back to width 512; the projections are trained for 20,000 updates to reconstruct the standardized features. Both quantized representations score below the continuous representation in all three categories, with homology falling from 72.65 to 62.62 with post-hoc VQ and to 28.40 with FSQ-4096.

##### Chain mixing

We assess the role of chain-context processing by removing the encoder’s bidirectional Mamba layers while retaining the GVP blocks and the two-layer Mamba predictor. Homology falls from 72.65 to 52.04; functional prediction also decreases (81.80 to 79.89), while flexibility is slightly higher (54.27 to 55.09). Because this removal also reduces encoder capacity, a parameter-matched variant expands the GVP blocks to a similar parameter count (14.057 versus 13.982 million). It recovers homology only partially (62.20), while its functional and flexibility scores exceed the reference (82.97 and 56.51). These results support a contribution of chain-context processing to homology prediction beyond parameter count, but not an advantage across all tasks.

##### Graph construction

With other settings fixed at 42,000 updates (Supplementary Section A.3), 30-neighbor graphs yield higher functional and flexibility scores and a similar homology score (72.83 versus 72.76) compared with 12-neighbor graphs.

Separately, at 35,000 updates, the C*α*-position and mean-position encoders score 81.35 versus 81.32, 53.46 versus 56.03, 72.93 versus 72.65 for function, flexibility and homology, respectively under masked latent prediction (Table S6(a,c)).

##### Pretraining method

We compare the principal PS-JEPA encoder used in Figure 2 with encoders pretrained by C*α* coordinate reconstruction (Table 1; encoder seeds 42–44, 35,000 updates each, original-label STB). The reconstruction encoder uses a C*α*-position graph and complete input structures, and is trained jointly with a two-layer bidirectional attention decoder (width 512, eight heads) that maps residue features to C*α* coordinates. The loss is the coordinate MSE after aligning each predicted chain to the reference with one translation and a proper rotation. STB uses the frozen non-EMA reconstruction encoder without the decoder, while PS-JEPA uses its frozen EMA encoder. Masking, graph positions, training schedules, and supervision scope also differ between these pretraining protocols.

##### Random encoder

Random encoders use PS-JEPA’s geometric inputs and architecture, remaining frozen during head training. Their mean functional, flexibility, and homology scores are 74.85, 44.27, and 27.14, respectively, below PS-JEPA (Table 1). Inputs and architectural biases provide predictive information before pretraining; PS-JEPA’s higher scores support an additional contribution from pretraining.

## B. StructTokenBench evaluation

### B.1. Evaluation protocol

STB (Yuan et al., 2025) comprises 12 tasks and 24 conditions on the released train/validation/test splits.

#### Heads

We add learnable positional embeddings to frozen encoder features and apply LayerNorm and dropout (0.1) before a two-layer MLP head. The MLP has a 512-unit ReLU hidden layer (128 in width-128 ablations). Homology classification uses masked mean pooling over valid residues to obtain a chain representation. Training follows Table S1, with head seed 1234 unless specified otherwise. We select head checkpoints by validation F1 for classification and validation Spearman correlation for regression, then evaluate the selected checkpoints on the test splits.

#### Metrics

Following the released STB implementation, functional AUROC and homology macro F1 use unweighted batch averages, including the final incomplete batch, with batch sizes of 8 and 64, respectively. Homology uses TorchMetrics’ multiclass macro F1, not pooled test-set F1. Flexibility Spearman uses concatenated valid predictions/targets. Category scores average over 15 functional, six flexibility, and three homology conditions. All benchmark scores are multiplied by 100 for readability.

#### Labels

Table S7 retains the original labels for comparability with the released STB benchmark. We also evaluated a corrected label version, corrected-v1, to assess the impact of identified label-mapping and target-precision errors. It aligns InterPro/epitope labels to sequence and BioLiP2 labels to residue identifiers and amino-acid identity; uncertain residues/samples are masked/excluded. ATLAS targets retain floating-point precision. Paired comparisons share samples/masks and refit heads. Splits and sequence overlaps are unchanged. Category-level performance changed little after label correction in the evaluated comparisons.

### B.2. Full results

We use released baseline weights or tokenizers, with adapters that align their outputs to residue positions. The heads receive fixed codebook embeddings or continuous residue features, depending on the representation. MiAE-S/B use unmasked backbone features; GearNet-Edge uses `mc_gearnet_edge`; ProteinAE uses eight-dimensional latents. MIF and GearNet also consume amino-acid identities (Table S7).

**Table S7:** StructTokenBench comparison of 13 frozen representations across 24 conditions in functional prediction, flexibility, and remote homology.

| Task | Split | Discrete / quantized |  |  |  |  |  | Continuous |  |  |  |  |  |  |
|---|---|---|---|---|---|---|---|---|---|---|---|---|---|---|
|  |  | Fold<br>seek | Pro<br>Tokens | ESM3<br>tokenizer | Vanilla<br>VQ | Amino<br>Aseed | Bio2<br>Token | Protein<br>MPNN | MIF<sup>†</sup> | MiAE<br>S | MiAE<br>B | Gear<br>Net<sup>†</sup> | Protein<br>AE | PS-JEPA<br>(*n* = 3) |
| **Functional prediction** *(AUROC)* |  |  |  |  |  |  |  |  |  |  |  |  |  |  |
| BindInt | Fold | 52.94<br>(53.18) | 43.86<br>(44.66) | 45.18<br>(44.30) | 47.74<br>(47.25) | 46.15<br>(47.11) | 45.07 | 52.13<br>(51.83) | 49.01<br>(50.38) | 55.27 | **58.32** | 53.34 | 45.57 | 50.95±0.87 |
|  | SupFam | 46.89<br>(46.20) | 86.45<br>(86.05) | 89.55<br>(90.77) | 86.87<br>(86.71) | 90.92<br>(90.53) | 84.23 | 94.00<br>(94.00) | 94.72<br>(94.56) | 95.87 | 96.14 | 96.09 | 82.25 | **97.25**±0.07 |
| BindBio | Fold | 52.23<br>(52.37) | 58.81<br>(58.47) | 63.20<br>(62.84) | 62.68<br>(62.02) | 65.66<br>(65.73) | 61.59 | 77.13<br>(78.42) | **86.14**<br>(85.79) | 77.90 | 80.00 | 82.85 | 67.01 | 79.75±0.92 |
|  | SupFam | 52.23<br>(52.41) | 60.82<br>(60.47) | 65.98<br>(65.22) | 63.52<br>(62.92) | 68.61<br>(68.30) | 63.19 | 79.13<br>(81.00) | 87.88<br>(87.27) | 80.46 | 83.75 | **88.02** | 67.73 | 85.78±0.27 |
| BindShake | Org | 53.23<br>(53.40) | 59.65<br>(59.82) | 66.51<br>(66.10) | 66.19<br>(67.04) | 68.30<br>(69.61) | 68.36 | 74.79<br>(75.52) | 80.98<br>(79.90) | 85.53 | **88.46** | 81.56 | 78.10 | 86.76±1.12 |
| CatInt | Fold | 53.32<br>(53.43) | 58.29<br>(58.16) | 60.12<br>(61.09) | 58.63<br>(58.89) | 61.90<br>(62.19) | 55.63 | 60.19<br>(61.05) | 60.34<br>(59.62) | 70.22 | 67.59 | 69.23 | 58.53 | **71.16**±2.71 |
|  | SupFam | 51.12<br>(51.41) | 84.03<br>(83.85) | 89.83<br>(89.82) | 85.09<br>(85.00) | 91.85<br>(91.91) | 82.02 | 93.45<br>(93.40) | 95.89<br>(96.49) | 96.19 | 97.09 | **97.71** | 79.33 | 97.66±0.11 |
| **Functional prediction (continued)** *(AUROC)* |  |  |  |  |  |  |  |  |  |  |  |  |  |  |
| CatBio | Fold | 55.03<br>(56.37) | 56.11<br>(56.14) | 65.25<br>(65.33) | 67.42<br>(67.58) | 65.35<br>(65.95) | 62.22 | 82.21<br>(82.49) | 84.94<br>(85.85) | 83.34 | 84.64 | **86.62** | 75.30 | 82.61±0.62 |
|  | SupFam | 53.73<br>(53.78) | 62.10<br>(64.05) | 74.68<br>(74.65) | 70.94<br>(70.92) | 87.65<br>(87.59) | 70.50 | 90.20<br>(93.19) | 96.95<br>(96.97) | 93.79 | 96.14 | **98.56** | 80.42 | 97.38±0.11 |
| Con | Fold | 49.30<br>(49.26) | 56.28<br>(56.23) | 55.47<br>(55.22) | 55.77<br>(56.98) | 57.31<br>(57.23) | 55.62 | 56.25<br>(57.18) | 60.10<br>(58.43) | 65.79 | **66.41** | 62.66 | 60.92 | 62.84±2.52 |
|  | SupFam | 51.33<br>(51.39) | 74.45<br>(74.33) | 79.31<br>(80.53) | 75.44<br>(74.60) | 86.63<br>(86.60) | 74.44 | 84.94<br>(84.68) | 91.22<br>(92.66) | 93.32 | 95.83 | 97.02 | 74.52 | **97.31**±0.07 |
| Rep | Fold | 47.12<br>(47.70) | 77.38<br>(77.25) | 75.62<br>(74.70) | 76.01<br>(75.99) | 75.49<br>(74.97) | 76.86 | **77.46**<br>(77.63) | 74.80<br>(74.53) | 75.03 | 74.63 | 74.22 | 74.81 | 75.04±0.73 |
|  | SupFam | 51.04<br>(52.53) | 78.80<br>(78.90) | 82.41<br>(82.36) | 82.14<br>(82.09) | 85.48<br>(84.57) | 79.13 | 81.01<br>(80.71) | 83.25<br>(83.11) | 80.39 | 82.88 | **88.72** | 79.64 | 86.49±1.04 |
| Ept | Fold | 52.78<br>(54.52) | 55.67<br>(54.69) | 60.84<br>(63.69) | 59.19<br>(59.28) | 62.70<br>(62.16) | 56.24 | 65.96<br>(62.84) | **70.41**<br>(68.78) | 64.14 | 62.81 | 69.98 | 51.94 | 67.46±1.69 |
|  | SupFam | 51.32<br>(50.56) | 68.03<br>(67.52) | 72.21<br>(61.97) | 67.18<br>(67.24) | 72.55<br>(72.02) | 69.69 | 77.47<br>(64.84) | 83.05<br>(82.98) | 72.66 | 76.52 | **83.21** | 61.04 | 82.25±0.38 |
| **Average** |  | 51.57 | 65.38 | 69.74 | 68.32 | 72.44 | 66.99 | 76.42 | 79.98 | 79.33 | 80.75 | 81.98 | 69.14 | 81.38 ± 0.14 |
| **Flexibility** *(Spearman)* |  |  |  |  |  |  |  |  |  |  |  |  |  |  |
| FlexRMSF | Fold | 14.80<br>(15.35) | 13.73<br>(13.81) | 43.71<br>(44.53) | 44.09<br>(44.22) | 44.54<br>(44.63) | 35.65 | 62.27<br>(62.37) | 59.87<br>(59.60) | 64.00 | **65.59** | 54.71 | 43.04 | 62.46±2.83 |
|  | SupFam | 11.61<br>(11.99) | 7.57<br>(7.62) | 40.87<br>(39.68) | 39.30<br>(39.08) | 40.67<br>(40.99) | 30.61 | 59.19<br>(59.24) | 56.73<br>(56.80) | 64.66 | **68.09** | 55.11 | 41.70 | 63.70±2.28 |
| FlexBFactor | Fold | 4.08<br>(4.17) | 7.62<br>(6.67) | 23.75<br>(23.60) | 22.36<br>(22.32) | 20.41<br>(21.30) | 20.05 | 32.21<br>(31.88) | **34.51**<br>(34.60) | 34.03 | 33.82 | 29.98 | 29.74 | 31.49±4.54 |
|  | SupFam | 6.89<br>(6.97) | 5.29<br>(5.47) | 25.77<br>(25.80) | 24.49<br>(23.73) | 21.43<br>(21.76) | 19.25 | 33.66<br>(34.56) | 35.32<br>(35.23) | 34.95 | 38.30 | 31.66 | 25.04 | **38.62**±1.89 |
| FlexNEQ | Fold | 5.12<br>(5.71) | 12.20<br>(12.98) | 45.04<br>(45.08) | 35.95<br>(35.95) | 49.94<br>(49.64) | 26.10 | **69.41**<br>(69.69) | 65.37<br>(65.32) | 54.54 | 57.39 | 62.95 | 16.07 | 63.57±0.38 |
|  | SupFam | 3.02<br>(2.60) | 11.61<br>(12.50) | 45.22<br>(45.43) | 35.70<br>(35.61) | 50.05<br>(50.15) | 26.09 | **68.55**<br>(68.69) | 64.82<br>(64.82) | 52.17 | 55.22 | 62.39 | 15.29 | 63.42±0.31 |
| **Average** |  | 7.59 | 9.67 | 37.39 | 33.65 | 37.84 | 26.29 | 54.22 | 52.77 | 50.73 | 53.07 | 49.47 | 28.48 | 53.88 ± 1.94 |
| **Remote homology** *(macro-F1)* |  |  |  |  |  |  |  |  |  |  |  |  |  |  |
| Homo | Fold | 11.40<br>(11.57) | 7.28<br>(5.84) | 28.23<br>(30.02) | 18.45<br>(18.17) | 28.97<br>(29.87) | 7.57 | 22.93<br>(25.66) | 22.31<br>(22.56) | 40.52 | 46.62 | 41.48 | 7.51 | **55.61**±2.95 |
|  | SupFam | 5.12<br>(4.67) | 4.85<br>(6.17) | 24.70<br>(24.89) | 19.77<br>(22.10) | 37.26<br>(38.38) | 4.99 | 28.05<br>(30.83) | 34.74<br>(33.86) | 46.32 | 60.93 | 68.02 | 4.66 | **70.60**±2.59 |
|  | Fam | 15.63<br>(15.30) | 14.55<br>(18.33) | 54.64<br>(54.42) | 43.65<br>(47.18) | 64.45<br>(69.78) | 19.32 | 61.77<br>(63.33) | 72.36<br>(74.22) | 82.09 | 85.84 | **91.87** | 14.24 | 90.51±0.55 |
| **Average** |  | 10.71 | 8.89 | 35.86 | 27.29 | 43.56 | 10.62 | 37.58 | 43.14 | 56.31 | 64.46 | 67.12 | 8.80 | 72.24 ± 1.45 |

Our results are shown in black; published STB scores appear in gray parentheses (Yuan et al., 2025). Bold indicates the highest displayed local score for each condition. Metrics (×100, higher is better): functional AUROC (area under the receiver operating characteristic curve), flexibility Spearman correlation, and homology macro F1. †: MIF and GearNet-Edge use structure and amino-acid identities. Bind: binding sites; Cat: catalytic sites; Int: InterPro; Bio: BioLiP2; Shake: ProteinShake; Con: conserved sites; Rep: repeats; Ept: epitopes; Homo: homology; SupFam/Fam: superfamily/family holdout; Org: original split. FlexRMSF: root-mean-square fluctuation; FlexBFactor: crystallographic B-factor; FlexNEQ: entropy-based backbone deformability. PS-JEPA: mean ± sample SD across encoder seeds 42–44, after averaging head seeds 1234, 2345, 3456 within each encoder.

### B.3. Homology metric aggregation

The STB evaluation averages macro F1 over test batches of up to 64 domains, so the score depends on batch composition. We therefore recompute homology macro F1 over each complete test split (pooled macro F1) from the same predictions as the batch-averaged scores, using the frozen encoders of the main STB comparison and validation-selected heads; no encoder or head is retrained, and no checkpoint is reselected. We compare PS-JEPA with GearNet-Edge, MiAE-B, and MiAE-S, the three highest-scoring homology baselines in Figure 2. Pooled macro F1 follows the released TorchMetrics convention of omitting classes absent from both targets and predictions; a second calculation averages over all 45 benchmark classes (Table S8).

PS-JEPA retains the highest category mean under pooled macro F1 (84.74±1.76), followed by MiAE-B (79.68), GearNet-Edge (77.80), and MiAE-S (74.29); the fixed-45-class calculation yields the same ordering. MiAE-B and GearNet-Edge exchange ranks relative to batch-averaged F1. After averaging the three heads within each encoder, all three PS-JEPA encoders score above every audited baseline mean under all three aggregations.

**Table S8.** Homology performance under batch-averaged and pooled macro F1.

| Model | Batch | Pooled macro F1 |  |  |  | Fixed-45 |
|---|---|---|---|---|---|---|
|  | mean | Fold | SupFam | Fam | mean | mean |
| PS-JEPA | 72.24 ± 1.45 | 64.22 ± 3.69 | 91.33 ± 1.41 | 98.67 ± 0.39 | 84.74 ± 1.76 | 75.87 ± 1.19 |
| GearNet-Edge | 68.10 ± 1.80 | 48.04 ± 1.13 | 87.13 ± 0.62 | 98.22 ± 0.52 | 77.80 ± 0.64 | 71.07 ± 0.53 |
| MiAE-B | 63.58 ± 3.06 | 55.61 ± 4.33 | 86.02 ± 2.39 | 97.41 ± 0.72 | 79.68 ± 2.23 | 71.84 ± 2.01 |
| MiAE-S | 57.78 ± 1.34 | 51.56 ± 4.79 | 78.12 ± 1.63 | 93.18 ± 1.33 | 74.29 ± 1.78 | 68.22 ± 1.14 |

Batch mean uses the original STB order, batch size 64, and an unweighted average including the final incomplete batch. Fixed-45 mean uses all 45 classes. PS-JEPA SD: three encoder means, each averaging three heads. Baseline SD: three heads for GearNet-Edge, MiAE-B, and MiAE-S, on one encoder per model. Figure 2 uses baseline head seed 1234.

## C. Backbone reconstruction

### C.1. Decoder and evaluation

To assess whether PS-JEPA representations retain recoverable backbone geometry without coordinate-reconstruction pretraining, we train a separate decoder while keeping the encoder frozen. Our conditional flow-matching decoder (Lipman et al., 2023) predicts the clean N/C*α*/C/O backbone from noisy coordinates, flow time, and frozen PS-JEPA embeddings *Z* ∈ ℝ^*L×*512^ of the complete input (Figure S4).

#### Architecture

Residue embeddings and the 12 N/C*α*/C/O coordinate components are separately projected to width 512 and added to learned positional and binary residue-mask embeddings. The decoder has 12 Transformer blocks (Vaswani et al., 2017), eight attention heads, and feed-forward width 2,048. An MLP maps 128-dimensional sinusoidal time features to DiT-style adaLN-Zero conditioning (Peebles and Xie, 2023).

The output head represents each residue using a rigid frame and a carbonyl rotation. These are parameterized by 11 values per residue: translation (3), a continuous 6D rotation representation (6), and a normalized sine–cosine pair for carbonyl rotation (2). The rotation obtained by Gram– Schmidt orthogonalization (Zhou et al., 2019) and the translation place an idealized N/C*α*/C template; the angle places O around the C*α*–C axis. Outputs are centered over observed atoms.

**Figure S4.**
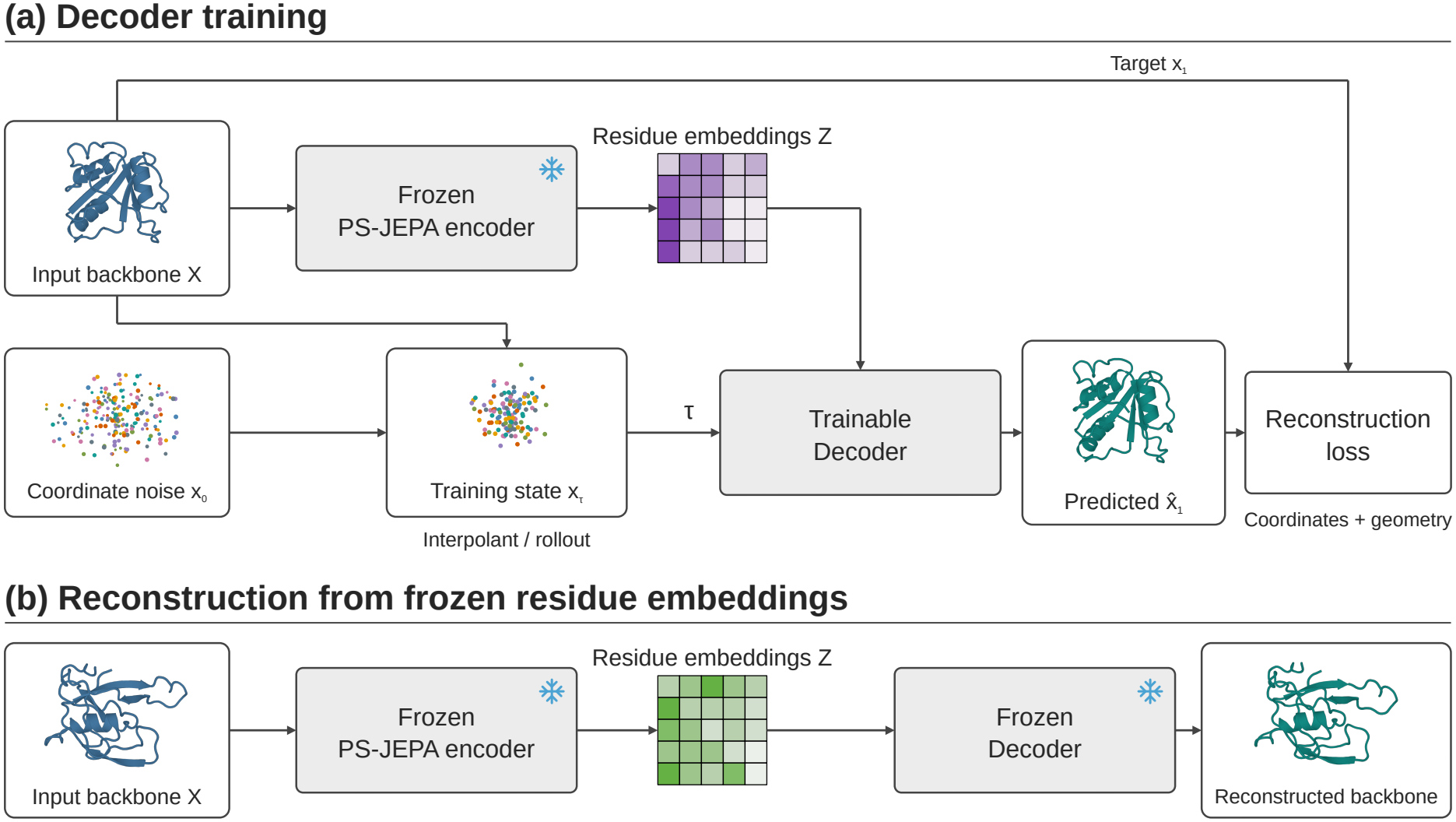
Conditional backbone reconstruction. (a) Frozen PS-JEPA embeddings condition decoder training using noise–target interpolants and rollout states. Only the decoder is updated. (b) With embeddings fixed, 100 Euler steps transform Gaussian noise into a reconstructed backbone. Structures illustrate two 120-residue AFDB training chains and their reconstructions.

#### Targets and initial objective

Coordinates are centered over observed atoms and divided by *s* = 10 Å; padding and unobserved atoms are masked throughout. A proper rotation, computed without gradients, aligns the target to centered Gaussian noise *x*_0_, giving *x*_1_. We use the interpolant *x*_*t*_ = (1 − *t*)*x*_0_ + *tx*_1_ and predict 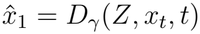. Initial training uses equally weighted coordinate MSE and unclamped frame-aligned point error (FAPE) (Jumper et al., 2021), both in scaled coordinates. MSE averages over valid atom-coordinate components across the minibatch. FAPE uses reference and predicted N/C*α*/C frames, requiring complete reference triplets; it averages valid frame–atom distances within each chain, then over chains with valid frames. Its distance calculation adds 10^*−*8^ under the square root.

#### Global-shape refinement

This stage adds two losses that supervise overall backbone geometry: C*α* squared error after rigid alignment with a proper rotation, and pair-distance MSE for residue pairs with |*i* − *j*| ≥ 24. The objective is

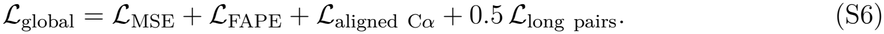

#### Rollout-based fine-tuning

The decoder is then trained on intermediate states generated by its own sampler. Each state is produced by eight Euler steps from independent Gaussian noise to a stopping time *t*^*^ drawn uniformly from {0.25, 0.5, 0.75, 0.99}, without interpolating toward target coordinates. For the rollout loss, gradients are computed only through the decoder’s prediction of the clean backbone from this state, not through the rollout or the rigid alignments between target and prediction. The objective is

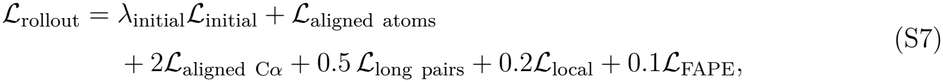

where *λ*_initial_ is the stage-specific weight of the initial MSE-plus-FAPE objective. *L*_initial_ is evaluated on an independently sampled noise–target interpolant; the remaining terms are evaluated on the prediction from the rollout state.

In both stages, the time of the noise–target interpolant follows 0.2*δ*_0_ + 0.4*U*(0, 0.2) + 0.4 *U* (0, 1) per chain, where *δ*_0_ denotes a point mass at zero. In rollout-based fine-tuning, this time is sampled independently of the stopping time *t*^*^. Aligned atom MSE averages over Cartesian components; aligned C*α* error sums over them. The local term is smooth L1 on C*α* pair distances below 15 Å in the reference, with a 1 Å transition width. All losses in both stages use scaled coordinates and average within chains before averaging across chains.

#### Training and checkpoint selection

We train the selected decoder on 742,183 AFDB chains of 18–512 residues in stages: 50k initial updates, 3k global-shape refinement updates, and three rollout stages of 3k, 12.5k, and 16k updates (84.5k total). We use *λ*_initial_ = 1*/*4, 1*/*8, 1*/*32 in the three rollout stages, respectively. Both stages use microbatches of four, four accumulation steps, and equal exposure to length bands 18–127, 128–255, 256–383, and 384–512. Each stage starts a new AdamW optimizer and cosine schedule (Supplementary Section A.1). BF16 mixed precision is used in both stages, with the frame projection and losses in FP32.

In both stages, checkpoints are evaluated every 500 updates on 128 fixed validation chains: 32 each from the 64–127, 128–255, 256–383, and 384–512 residue bands. Global-shape refinement selects the checkpoint with the lowest mean C*α* RMSD. Rollout stages minimize population-weighted C*α* RMSD, using the length-band frequencies in the original 6,956-chain development pool as weights. Relative to the checkpoint at the start of each rollout stage, eligible checkpoints must not decrease population-weighted C*α*-lDDT by more than 0.002 and must keep each band’s C*α* RMSD at or below 1.03× its starting value + 0.01 Å. This selection cohort is reused across experiments.

#### Sampling

Holding *Z* fixed, we take 100 Euler steps from centered Gaussian noise at *t*_*k*_ = *k/*100, *k* = 0, …, 99, with velocity (*D*_*γ*_(*Z, x*_*t*_, *t*) − *x*_*t*_)*/*(1 − *t*). Each step recenters valid atoms; final coordinates are multiplied by *s* to restore Å. Reference coordinates enter through *Z* and subsequent evaluation, not the sampled coordinate state. Unconditional backbone generation uses this same checkpoint with decoder weights frozen (Supplementary Section D.1).

#### Reconstruction metrics

Comparisons use observed-atom masks and fixed residue correspondence; cohort summaries weight chains equally. C*α* and backbone RMSD use separate proper-rotation Kabsch fits (Kabsch, 1976) over observed C*α* and N/C*α*/C/O atoms, respectively. The main text reports C*α* RMSD. TM-score (Zhang and Skolnick, 2004) uses US-align (Zhang et al., 2022) (-TMscore 1) with residue-index correspondence, optimizing superposition and normalizing by the observed reference C*α* count. Our pair-weighted C*α*-lDDT (Mariani et al., 2013) pools distinct unordered observed C*α* pairs with reference distance below 15 Å and averages the fractions with absolute distance error strictly below 0.5, 1, 2, and 4 Å. Pairs are weighted equally within each chain without first averaging per-residue scores; adjacent residues are included, self-pairs excluded, and no stereochemical penalty applied. RMSD uses unrounded inference and reference coordinates; TM-score uses both structures saved to PDB at 0.001 Å precision; C*α*-lDDT uses the saved prediction and unrounded reference coordinates.

#### External reconstruction test

Both checkpoints and inference settings are fixed before external evaluation. We query all 2026 PDB releases through 12 November for X-ray protein entities of 64–512 residues at resolution ≤ 3 Å (Berman et al., 2000). Entries identified in pre-existing local structure files and caches are excluded. Among 9,030 entities with canonical sequences, MMseqs2 (Steinegger and Söding, 2017) excludes alignments with identity ≥ 30% covering ≥ 80% of the query against AFDB training/validation, cached PDB reference sets, and prior STB evaluation sequences, leaving 983 entities. Within each entity, we use the first lexicographically ordered chain meeting 64–512 observed residues and ≥ 90% C*α* coverage. After this quality check, deterministic greedy selection removes within-test matches at ≥ 30% identity and ≥ 80% coverage of both sequences. The resulting 309-chain manifest is defined without using reconstruction scores. Each chain is reconstructed once using a preassigned noise seed and the same 100-step sampler and metrics described above; all declared test chains are included. Foldseek (van Kempen et al., 2024) additionally searches AFDB training structures. The TM-filtered sensitivity subset removes matches with both query- and target-normalized TM-scores ≥ 0.5. These finite-sensitivity searches do not establish absence of all remote homology; the primary external cohort is not claimed to be fold-disjoint.

## D. Unconditional generation

### D.1. Prior training and sampling

Our length-conditioned flow-matching prior (Lipman et al., 2023) models standardized residue features *Y* = (*Z* − *µ*)*/σ*, where *Z* = *E*(*X*) ∈ ℝ^*L×*512^ denotes frozen teacher encoder embeddings and *µ* and *σ* are the channel means and standard deviations of the filtered AFDB training pool. After confidence filtering (mean pLDDT ≥ 85 and pLDDT standard deviation ≤ 15, using the released Proteina *D*_21M_ index (Geffner et al., 2025)) and backbone-geometry filtering (complete N/C*α*/C/O coordinates, adjacent C–N distances of 1–2 Å, a C*α* radius of gyration no greater than 30 Å, and a DSSP (Kabsch and Sander, 1983) coil fraction no greater than 0.5), we retain 50,653 training chains and 528 validation chains of 64–128 residues. We train a DiT-style prior (Peebles and Xie, 2023) from random initialization, with the encoder and reconstruction decoder frozen; Table S9 gives the architecture and training settings.

Given a Gaussian noise–target interpolant *Y*_*t*_ = (1 *− t*)*ϵ* + *tY*, the prior predicts the clean features *Ŷ* and minimizes a time-weighted MSE between *Ŷ* and *Y*. Every fourth update, we add a geometry loss (weight up to 0.05): the predicted features are decoded through eight differentiable Euler steps of the frozen reconstruction decoder, and coordinate, local-distance, and FAPE terms are computed against the training backbone (Supplementary Section C.1); gradients pass through the decoder to update the prior. We retain the checkpoint with the lowest validation latent loss on a fixed subset of 128 validation chains (update 41,000); this loss excludes the geometry loss. For sampling, we take 200 prior Euler steps and decode the sampled features using 100 Euler steps of the same frozen decoder (Figure S5).

**Table S9.**
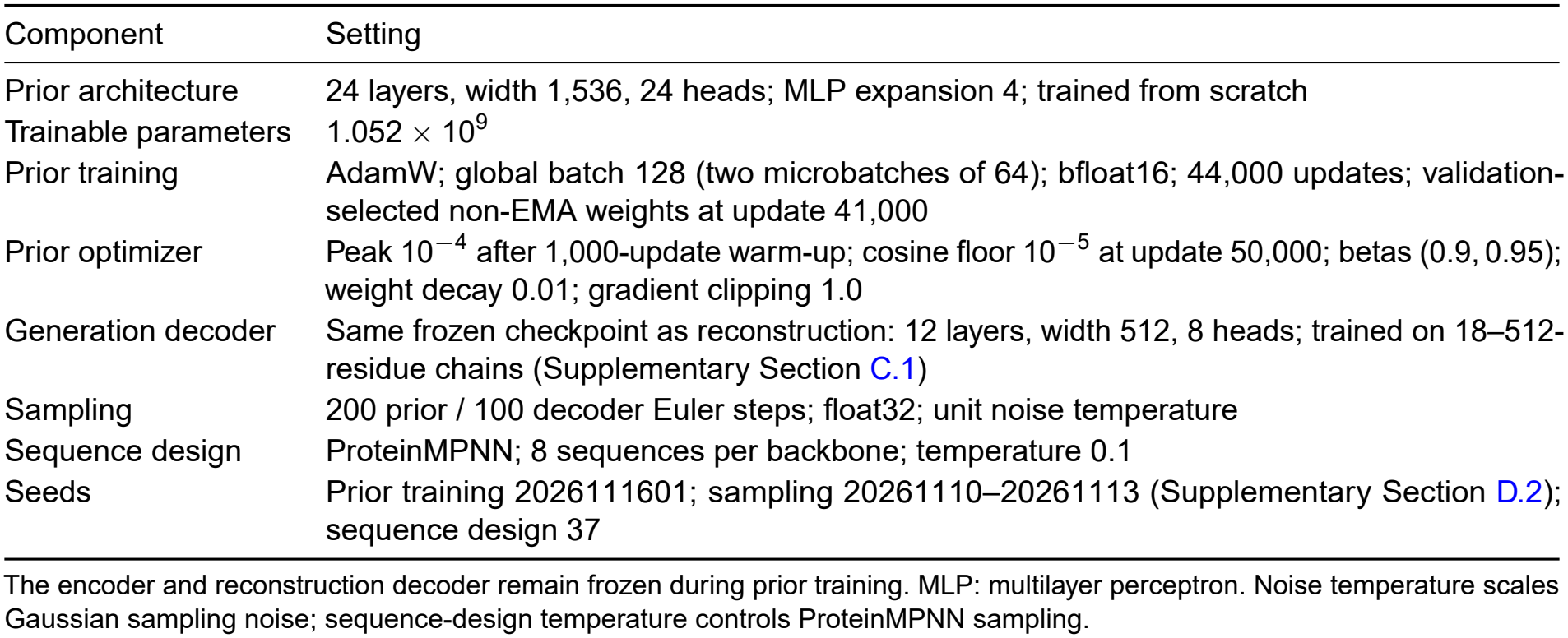
Training and sampling settings for the latent prior.

**Figure S5.**
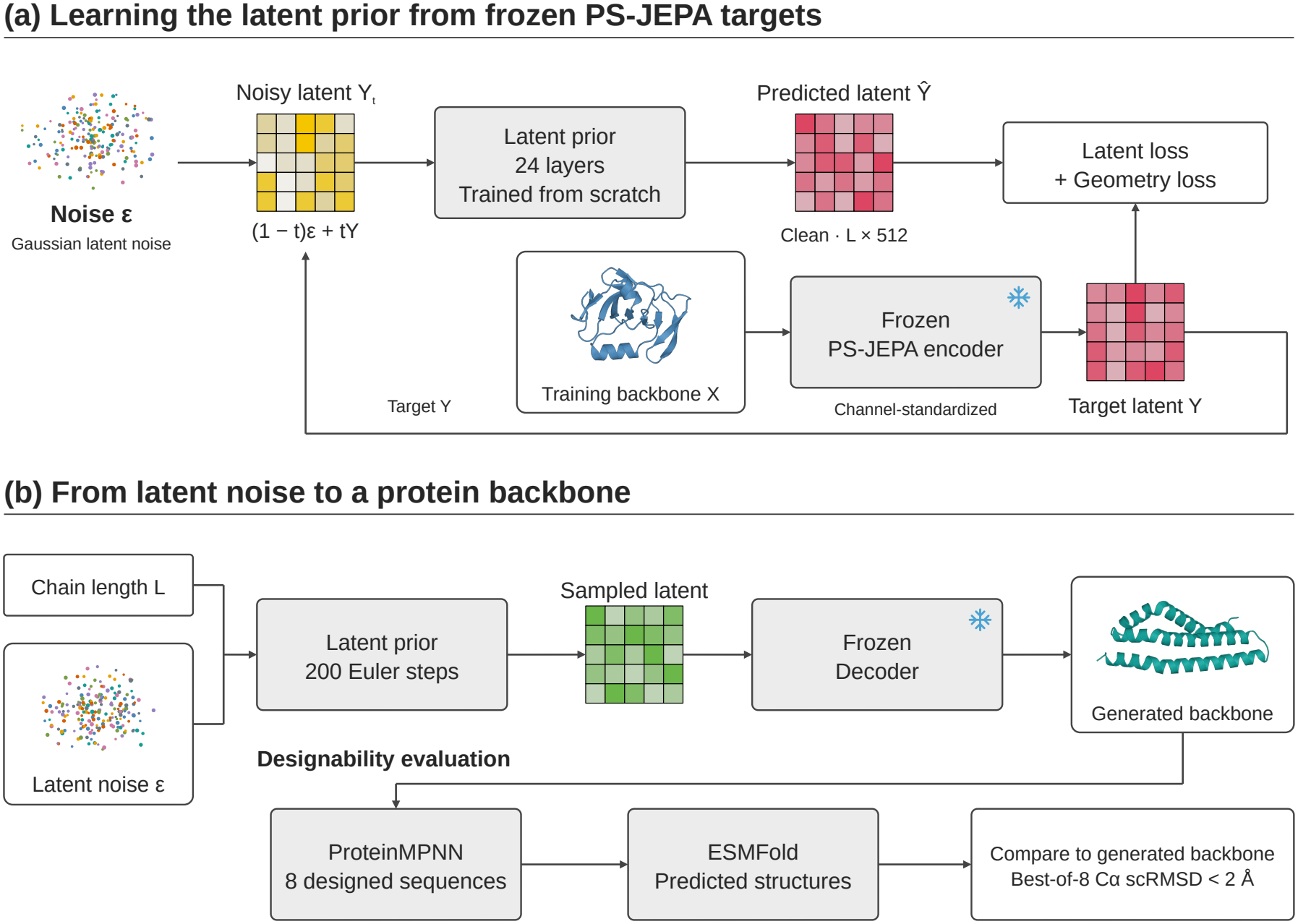
Unconditional backbone generation in frozen representation space. (a) Latent prediction and decoded-geometry losses train the prior from scratch; the encoder and decoder remain frozen. The geometry-loss path through the frozen decoder is omitted for clarity. (b) The prior is conditioned only on chain length and generates embeddings in 200 Euler steps, followed by 100 decoder steps to generate a backbone. ProteinMPNN designs eight sequences per backbone, which are refolded with ESMFold. A backbone is designable if at least one refold achieves full-chain C*α* scRMSD below 2 Å. The illustrated output is a 120-residue backbone from the evaluated cohort.

### D.2. Designability

We sample 32 backbones for each *L* ∈ {80, 100, 120} and seed 20261110–20261113, giving 128 per length and 384 total, without post-generation filtering. Each backbone receives eight ProteinMPNN-designed sequences refolded with ESMFold (Dauparas et al., 2022; Lin et al., 2023b), using the settings in Table S9. A backbone is classified as designable if at least one of its eight designed sequences refolds with full-chain C*α* scRMSD below 2 Å after proper-rotation alignment (Geffner et al., 2025). All 384 backbones remain in the denominator, including failed attempts. The four seeds vary sampling with the trained prior fixed. Structural novelty of the generated backbones relative to the training set of the prior was not assessed.

